# Assessment of CSF biomarkers and microRNA-mediated disease mechanisms in SMA

**DOI:** 10.1101/2021.05.19.444816

**Authors:** Emily Welby, Rebecca J. Rehborg, Matthew Harmelink, Allison D. Ebert

## Abstract

**Objective:** Cerebral spinal fluid (CSF) is a promising biospecimen for the detection of central nervous system (CNS) biomarkers to monitor therapeutic efficacy at the cellular level in neurological diseases. Spinal muscular atrophy (SMA) patients receiving intrathecal antisense oligonucleotide (nusinersen) therapy tend to show improved motor function, but the treatment effect on cellular function remains unknown. The objective of this study was to assess the potential of extracellular RNAs and microRNAs in SMA patient CSF as indicators of neuron and glial health following nusinersen treatment.

**Methods:** CSF samples from SMA Type 1-3 patients were screened using quantitative RT-PCR to assess expression of extracellular RNAs associated with inflammation and cellular stress, and microRNAs previously implicated in SMA pathogenesis. We also used mRNA sequencing and multi-electrode array approaches to assess the transcriptional and functional effects of astrocyte-associated miR-146a on healthy and SMA induced pluripotent stem cell (iPSC)-derived motor neurons.

**Results:** Extracellular RNA analysis is suggestive of ongoing cellular stress, even in nusinersen treated samples. microRNAs previously associated with SMA pathology tended to show improvement in expression levels in nusinersen treated samples, with the exception of the astrocyte-secreted miR-146a. miR-146a treated iPSC-derived motor neurons showed a downregulation of extracellular matrix genes found in the synaptic perineuronal net and decreased spontaneous activity.

**Interpretation:** Extracellular RNAs and microRNAs can be detected in SMA patient CSF samples, potentially serving as useful biomarkers to monitor cellular health during nusinersen treatment. Astrocyte health and response to nusinersen are important aspects to address in SMA pathogenesis and treatment strategies.

## Introduction

Spinal muscular atrophy (SMA) is the leading genetic cause of pediatric mortality (1) and is caused by the homozygous disruption of the *Survival of Motor Neuron-1* (*SMN1*) gene. *SMN1* is responsible for generating full length SMN protein, which has an established global role in small nuclear ribonucleoprotein (RNP) assembly and pre-mRNA splicing (2), and is particularly important for motor neuron function (3, 4). Reduced SMN leads to selective lower spinal motor neuron degeneration and subsequent loss of skeletal muscle innervation leading to paralysis, respiratory failure, and premature death. A human specific *SMN1* paralog, *SMN2*, can partially produce full length SMN protein and modify SMA disease severity (5, 6), but a single nucleotide change within the splicing enhancer region of *SMN2* exon 7 (c.840C>T) causes aberrant exon 7 splicing, generating an unstable, non-functional SMN protein which is targeted for degradation (7, 8).

Efforts to develop antisense oligonucleotide technology to facilitate *SMN2* splicing correction and increase full length SMN protein levels led to the first FDA approved drug, nusinersen, for SMA (9). Nusinersen is delivered intrathecally into the CNS, where it targets the *SMN2* intronic splicing silencer, blocking the binding of heterogenous nuclear RNP A1 that represses exon 7 inclusion (10). Target specificity, minimal toxicity, and long half-life properties have been demonstrated for nusinersen and remarkable improvements in motor milestone achievements are observed in patients treated pre-symptomatically (11, 12). However, improvements at the cellular level post treatment is not fully understood. CSF biomarker detection that can reliably track improvements in CNS cellular health and correlate with motor functioning recovery has high clinical value. In this regard, studies have begun to assess biomarkers in SMA CSF samples after nusinersen treatment, such as neurofilament light chain (13, 14) and synaptic-associated proteins (15), however, the reliability of these markers is still in question. In other neurodegenerative diseases, microRNAs are promising biomarkers detectable in CSF samples (16-18), but they have yet to be explored in SMA patient CSF samples. Further work is also needed to fully define SMA cell autonomous and non-cell autonomous disease mechanisms, which will ultimately impact biomarker discovery efforts.

In this study, we screened SMA patient CSF samples before and after multiple doses of nusinersen for extracellular RNAs and microRNAs to facilitate biomarker discovery. Overall, we observe that cellular stress RNAs are still present in the CSF even after treatment administration. Although we observe an overall post treatment improvement in microRNA expression levels previously associated with SMA disease pathology, the astrocyte-secreted miR-146a, which we previously show to be toxic to SMA motor neurons (19), remains elevated in SMA patient CSF. We further demonstrate miR-146a alters perineuronal net extracellular matrix gene expression in iPSC-derived motor neurons, which may affect synaptic functioning. Taken together, these data suggest that extracellular RNAs and microRNAs detected in patient CSF may be valuable for tracking cellular health in response to nusinersen treatment. Additional studies focused on astrocyte-mediated disease mechanisms, nusinersen uptake and cellular recovery will be important to ensure long term efficacy of therapies for SMA.

## Methods

### Human samples

CSF samples were obtained at Children’s Wisconsin with informed consent during standard clinical care for the administration of SMA-related nusinersen ASO therapy (IRB 1080061). A total of 35 CSF samples were used in this study from SMA Type 1 (4 patients), Type 2 (5 patients) and Type 3 (3 patients). Deidentified SMA patient information related to pre- and post-nusinersen treated CSF biospecimens analyzed in this study and motor functioning scores, are detailed in Table 1. Before each dose, the patients received a physical exam by a pediatric neuromuscular physician or physician’s assistant as well as complete blood count, urinalysis, complete metabolic panel, INR, PT and PTT. Additionally, there were periodic standardized assessments by a trained research physical therapist as required based upon the age of the patient, insurance payor needs for coverage of the drug, as well as patient availability. The use of CSF samples and iPSCs was approved by the Medical College of Wisconsin Institutional Review Board (PRO00030075; PRO00025822), the Institutional Biosafety Committee (IBC20120742) and the Human Stem Cell Research Oversight Committee.

**Table 1.** List of patient samples used in this study and motor function scores. CHOP-I-SCORE: The Children’s Hospital of Philadelphia Infant Test of Neuromuscular Disorders; HINE: Hammersmith Infant Neurological Examination; HFMSE: Hammersmith Functional Motor Scale Expanded; RULM: Revised Upper Limb Module.

| Patient | SMA Type | Gender | Age | Samples | Motor function scores |  |  |  |  |  |  |  |  |  |
| --- | --- | --- | --- | --- | --- | --- | --- | --- | --- | --- | --- | --- | --- | --- |
|  |  |  |  |  | Scores nearest CSF1 | Scores nearest CSF4 | Scores nearest CSF5 | Scores nearest CSF6 | Scores nearest CSF8 | Scores nearest CSF9 | Scores nearest CSF10 | Scores nearest CSF11 | Scores nearest CSF12 | Scores nearest CSF13 |
| SMA006 | 1 | Female | 7 months | CSF 1, 4 | CHOP-I-SCORE:<br>22/64<br>HINE<br>SCORE:<br>27/78 |  |  |  |  |  |  |  |  |  |
| SMA017 | 1 | Female | 2 months | CSF 1, 4, 5 | CHOP-I-SCORE:<br>10/64<br>HINE<br>SCORE:<br>28/78 |  | CHOP-I-SCORE:<br>26/64<br>HINE<br>SCORE:<br>35/78 |  |  |  |  |  |  |  |
| SMA001 | 1 | Female | 9 months | CSF 1, 4, 8 | CHOP-I-SCORE:<br>22/64<br>HINE<br>SCORE:<br>29/78 |  |  |  | CHOP-I-SCORE:<br>37/64<br>HINE<br>SCORE:<br>29/78 |  |  |  |  |  |
| SMA021 | 1 | Female | 20yrs | CSF 1, 4, 5 | Physical<br>therapy<br>tests not<br>obtained |  |  |  |  |  |  |  |  |  |
| SMA002 | 2 | Female | 3yrs | CSF 1, 4, 13 | HFMSE<br>SCORE:<br>3/66<br>RULM<br>SCORE:<br>10 | HFMSE<br>SCORE:<br>4/66<br>RULM<br>SCORE:<br>12 |  |  |  |  |  |  |  | HFMSE<br>SCORE:<br>10/66<br>RULM<br>SCORE:<br>14 |
| SMA003 | 2 | Female | 11yr | CSF 1, 4, 13 | HFMSE<br>SCORE:<br>10/66<br>RULM<br>SCORE:<br>19/38 | HFMSE<br>SCORE:<br>10/66<br>RULM<br>SCORE:<br>21 |  |  |  |  |  |  |  | HFMSE<br>SCORE:<br>6/66<br>RULM<br>SCORE:<br>14 |
| SMA022 | 2 | Male | 11yrs | CSF 1, 4, 6 | HFMSE<br>SCORE:<br>6/66<br>RULM<br>SCORE:<br>20 |  |  |  |  |  |  |  |  |  |
| SMA018 | 2 | Female | 3yrs | CSF 1, 4, 10 | CHOP-I-SCORE:<br>38/64<br>HINE<br>SCORE:<br>38/78<br>HFMSE<br>SCORE:<br>11/66 | CHOP-I-SCORE:<br>43/64<br>HFMSE<br>SCORE:<br>14/66<br>RULM<br>SCORE:<br>14 |  |  |  |  | HFMSE<br>SCORE:<br>15/66<br>RULM<br>SCORE:<br>15 |  |  |  |
| SMA010 | 2 | Male | 13yrs | CSF 1, 4, 11 | HFMSE<br>SCORE:<br>2/66<br>RULM<br>SCORE:<br>6 | HFMSE<br>SCORE:<br>2/66<br>RULM<br>SCORE: 8 |  |  |  |  |  | HFMSE<br>SCORE:<br>2/66<br>RULM<br>SCORE:<br>7 |  |  |
| SMA007 | 3 | Male | 14yrs | CSF 1, 4, 9 | HFMSE<br>SCORE:<br>51/66<br>RULM<br>SCORE:<br>22 | HFMSE<br>SCORE:<br>46/66<br>RULM<br>SCORE:<br>23 |  |  |  | HFMSE<br>SCORE:<br>52/66<br>RULM<br>SCORE:<br>27 |  |  |  |  |
| SMA023 | 3 | Male | 4yrs | CSF 1, 4, 6 | HFMSE<br>SCORE:<br>49/66<br>RULM<br>SCORE:<br>28/37 | HFMSE<br>SCORE:<br>52/66<br>RULM<br>SCORE:<br>32 |  |  |  |  |  |  |  |  |
| SMA004 | 3 | Male | 9yrs | CSF 1, 4, 12 | HFMSE<br>SCORE:<br>39/66<br>RULM<br>SCORE:<br>27 | HFMSE<br>SCORE:<br>40/66<br>RULM<br>SCORE:<br>27 |  |  |  |  |  |  | HFMSE<br>SCORE:<br>39/66<br>RULM<br>SCORE:<br>32 |  |

### RNA extraction and spike-in control assessment from patient CSF samples

CSF samples were immediately frozen after collection and stored at −80°C. CSF samples visibly contaminated with blood were not used in this study. On day of processing, CSF were thawed on ice, centrifuged for 5 minutes at 700 x g to remove any contaminating debris, and aliquoted into fresh tubes. 1x 625ul aliquot of each CSF sample was processed immediately for RNA extraction using the mirVana™ PARIS™ RNA and Native Protein Purification Kit (Ambion), following the liquid sample preparation according to manufacturer’s instructions. Before the addition of the denaturing solution, 1ug of MS2 RNA carrier (Sigma) was added to each sample to facilitate RNA recovery. Prior to acid-phenol:chloroform addition, 3.5ul of miRNeasy Serum/Plasma Spike-In Control (miR-39, 1.6 × 10^8^ copies; Qiagen) was added to each CSF sample. qPCR to detect this miR-39 spike-in control was performed post extraction to estimate RNA recovery. cDNA was generated from CSF isolated RNA and a control RNA sample spiked with miR-39 (1 × 10^8^ copies/ul) using the miScript HiSpec kit (Qiagen); dilutions were created from the control cDNA sample spiked with miR-39 (1 ×10^6^-1×10^3^ copies) to generate a standard curve. The miR-39 spike-in control copies in CSF samples ranged from 40,257-986,469 copies; these data were used to normalize RNA input across CSF samples for RT-qPCR experiments.

### RT-qPCR array

RNA samples were DNase treated using the TURBO DNA free kit (Ambion) prior to cDNA synthesis. The RT^2^ PreAMP cDNA synthesis kit (Qiagen) was used with the Human Aging RT^2^ PreAMP Pathway Primer Mix. Samples were applied to the Human Aging RT^2^ Profiler PCR arrays (Qiagen) and run on Bio-Rad CFX384 real time thermocycler. Samples passed quality controls built into the array plates and target gene Cq values on qPCR array were normalized to the HPRT1 house keeping gene values. Ct values observed in the latest post-treatment CSF samples collected at the time of this study for each patient were normalized to pre-treatment (CSF1) samples to calculate the relative fold change (ddCT) in gene expression.

### microRNA qPCRs

cDNA for microRNA qPCR experiments was generated from CSF RNA samples using the miScript II RT kit. Preamplification on cDNA was performed using the miScript PreAMP kit using the following primer sets (Qiagen): Hs miR146a_1 (M00003535), Hs miR23a_2 (M00031633), Hs_mir-218_1 (MS00006769), Hs_mir-9_1 (MS00010752), Hs_mir-132_1 (MS00003458) and Hs_mir-431_1 (MS00004200). Hs miR103a_1 (M00031241) was included to be used as a housekeeping microRNA for qPCRs. PreAMP primer sets for miR-16 (determining dilution factor) and mirTC (estimated efficiency of preamplification reactions) were also included in the amplification reactions; CSF samples did not require further dilution and all samples passed the preamplification quality control. qPCRs for each target microRNA was performed using miScript SYBR Green PCR kit (Qiagen) using a Bio-Rad CFX384 real time thermocycler. In this analysis, we used CSF1 (pre-treatment), CSF4 (4^th^ nusinersen dose) and the latest post-treatment CSF samples collected at the time of this study for each patient. All samples were normalized to pre-treatment samples (CSF1) for each patient to calculate the relative fold change in expression.

### Stem cell culture, motor neuron differentiation and mir146a mimic treatment

Previously characterized induced pluripotent stem cell lines (iPSCs) from 2x SMA Type 1 patients (7.12 and 3.6) and 2x healthy individuals (21.8, 4.2; 4.2 is the parent of 3.6) were used in this study (19, 20). iPSCs maintenance cultures, motor neuron differentiation and microRNA treatments were performed as previously described (19, 21). Briefly, on Day 28 of motor neuron differentiation, cells were treated for 48 hours with 40nM miR-146a-5p mimic (Ambion) or mirVana miRNA mimic negative control 1 (Ambion). EndoPorter (Gene Tools, LLC) was used as the transfection reagent. Cells were collected for RNA sequencing or analyzed in multi-electrode array (MEA) experiments.

### RNA sequencing of motor neuron differentiations

Cell pellets were processed using the RNeasy kit (Qiagen) according to manufacturer’s guidelines; on-column DNaseI treatment (Qiagen) was also performed on all samples. RNA concentrations and RIN values were obtained using the RNA ScreenTape kit and the Tapestation system (Agilent); all samples had a RIN score >9 with clear 18S and 28S peaks. ERCC Spike-In Controls (Invitrogen) were added to RNA samples (input RNA was 100ng per sample) prior to processing sample with the NEBNext Poly(A) mRNA Magnetic Isolation Module (NEB #E7490), NEBNext Ultra II RNA Library Prep Kit for Illumina (NEB # E7770) according to manufacturer’s protocol. NEBNext Multiple Oligos for Illumina Set 1 (NEB #E7335), 2 (NEB #E7500) and 3 (NEB #E7710) were used to create unique identifiers for each cDNA library sample. Quality of cDNA libraries were assessed using the DNA 100 Chip kit and Tapestation system (Agilent). Single and pooled libraries was assessed using the NEBNext Library Quant kit for Illumina (NEB #E7630), Bio-Rad CFX384 real time thermocycler and calculated using the NEB qPCR webtool (https://nebiocalculator.neb.com/qPCR). On the day of sequencing, samples were denatured and diluted following the Illumina NextSeq system Denature and Dilute Libraries Guide using the bead-based normalization method with PhiX library control. Samples were sequenced (20-30 million read depth, 38bp paired end reads) on the Illumina NextSeq system. FASTQ files were processed using Expression count (STAR) pipeline in Basepair before performing principal component analysis and DeSeq2 RNA seq analysis. Gene Ontology analysis was performed using the gprofiler web server (22).

### Multi-electrode array

On day 14 of motor neuron differentiation, neurospheres were dissociated in TrypLE (Life Technologies) and plated on to 0.1% PEI coated CytoView MEA 48 well plates (Axion Biosystems) at 50,000 cells per well in a 10ul droplet over the electrodes. 20µg/ml laminin was supplemented into the motor neuron maturation medium (composition described in (19, 21)) to aid cell attachment. After 1-hour incubation, wells were flooded with 300ul motor neuron maturation medium; medium was replaced every 3 days. Spontaneous activity recordings were performed using the Axion Biosystems Maestro System on a stage maintained at 37°C. 1-minute recordings were taken before (Day 28) and after (Day 30) treatment. Weighted mean firing rate (WMFR; number of spikes (i.e. action potentials) from active electrodes (>5 spikes) over the duration of analysis) measured in Hz was used to assess motor neuron activity in this study. Recordings, data collection and waveform analyses were completed using Axis (Axion Biosystems). WMFR treatment data was normalized to UTX control for each line.

### Statistics, rigor and reproducibility

For the qPCR array, 6x Type 1, 6x Type 2 and 2x Type 3 SMA patient samples were analyzed; for the microRNA qPCRs, 11x Type 1, 15x Type 2, 9x Type 3 SMA patient samples were analyzed. Samples were run in technical duplicates and Ct values were averaged for downstream analysis. For the RNA seq experiments, motor neurons were generated from 2x SMA Type 1 patients and 2x healthy individuals; 2-3 individual differentiation samples were included in the analysis. -log10 adjusted p-values (<0.05) and log2 fold change (<-1 or >1) thresholds were considered statistically significant for the DeSeq2 analysis. The same 4 cell lines were run in technical triplicates on the MEA plates; data are from 2x plates.

## Results

Surplus CSF was used in this study collected from SMA patients receiving intrathecal nusinersen injections during the course of standard clinical care. Table 1 details the patients’ gender, age, CSF samples analyzed in this study and motor functioning test scores. Clinically, all patients in this study receiving treatment did well without decline in feeding, respiratory support requirements or motor activities of daily living. One patient, SMA006, did transfer care to another center resulting in a lack of further assessments, while patient SMA001 died suddenly in her sleep without an overt decline or infection noted prior; no adverse effects from the medication were noted. Regarding the standardized assessments, while they did not correlate directly to the therapeutic infusions temporally, they did demonstrate an overall improvement in patient motor functioning. Patient SMA003 was an exception; the patient’s strength was improving but overall, there was a decline in the Hammersmith score secondary to the patient developing contractures from a lack of reliable stretching.

We used a quantitative RT-PCR (qRT-PCR) array approach to begin assessing novel extracellular mRNA biomarkers associated with inflammatory and stressor responses in SMA patient CSF samples. Across SMA Type 1 (Fig. 1Ai), Type 2 (Fig. 1Bi) and Type 3 (Table 2) patient samples, we identified a total of 48 upregulated genes, involved in immune responses (FCER1G, CD14), inflammatory signaling (C3AR1, TOLLIP, S100A8, S100A9), cell cycle regulation (CDKN1C, RAP1A), genomic stability (SIRT6) and apoptosis (CASP1, CLU, PDCD6). Furthermore, genes encoding fibril-forming collagen proteins (COL1A1 and COL3A1) were also found to be upregulated in Type 1 and 2 patients, while the TGF-β signal transducer, SMAD2, was found to be upregulated in both Type 2 and 3 patients. 46 downregulated genes were identified across all SMA disease types in samples post-nusinersen treatment (Fig 1A-Bii; Table 2); intriguingly, ANGEL2, a 2’,3’-cyclic phosphatase involved in RNA processing and unfolded protein response (UPR) transcription factor, XBP1, mRNA splicing (23), was the only gene to show consistent downregulation across all SMA disease types. ELAVL1, a known binding partner of SMN, also showed a downregulated expression trend, but only in SMA Type 1 patient samples. We did note some overlapping genes between the upregulated and downregulated datasets, dependent upon the SMA disease type. Gene Ontology (GO) analysis revealed an overall enrichment of proteolysis and metabolic processes terms within the upregulated genes (Fig. 1C), whereas astrocyte and glial cell development GO terms were particularly enriched within the downregulated genes (Fig. 1D).

**Table 2.** Gene lists of upregulated and downregulated genes from RT-qPCR array.

| Upregulated genes from qPCR array |  |  |  |
| --- | --- | --- | --- |
| Patient | Gene | CSF samples |  |
|  |  | Fold change (CSF1 vs CSF8) | Standard deviation |
| SMA Type 1 Patient 1 | CASP1 | 137.6780611 | 0.077066231 |
|  | RAP1A | 105.0277355 | 0.396821273 |
|  | FCER1G | 89.4378156 | 0.01024327 |
|  | SMAD2 | 75.600257 | 1.072265243 |
|  | S100A8 | 61.92780669 | 0.345332247 |
|  | CLU | 51.10488666 | 0.242735864 |
|  | TINF2 | 31.1294647 | 0.385072282 |
|  | COL3A1 | 4.993260377 | 0.567399927 |
|  |  | Fold change (CSF1 vs CSF4) | Standard deviation |
| SMA Type 1 Patient 6 | CDKN1C | 624.5714924 | 0.113307 |
|  | FCGR1A | 114.1071797 | 0.353113154 |
|  | LYZ | 109.0805366 | 0.17213043 |
|  | FCER1G | 106.5105179 | 0.078189651 |
|  | CD14 | 63.99338544 | 0.626886034 |
|  | TMEM135 | 54.11798305 | 1.054954958 |
|  | ELP3 | 46.86267237 | 1.866680798 |
|  | C3AR1 | 18.75636511 | 0.27921538 |
|  | PDCD6 | 3.179984491 | 0.273234679 |
|  | COL1A1 | 2.791978761 | 0.141905159 |
|  | ANXA5 | 1.80016017 | 0.178070963 |
|  |  | Fold change (CSF1 vs CSF5) | Standard deviation |
| SMA Type 1 Patient 17 | CD14 | 179.8169828 | 0.329436003 |
|  | TOLLIP | 92.04404226 | 0.720422074 |
|  | MBP | 89.23893429 | 0.155628565 |
|  | TPP1 | 86.18803061 | 0.116520265 |
|  | CDKN1C | 82.01489092 | 0.562600756 |
|  | EML1 | 81.69333115 | 0.271808028 |
|  | VPS13C | 68.11989442 | 0.352171864 |
|  | CASP1 | 66.34249587 | 0.578650204 |
|  | HSF1 | 52.15679859 | 0.044524139 |
|  | CLU | 37.00182299 | 0.02191813 |
|  | POT1 | 33.94264604 | 0.360925318 |
|  | PHF3 | 32.10791118 | 1.07868608 |
|  | MRPL43 | 21.92243282 | 0.29517183 |
|  | GSTA1 | 20.36415497 | 0.278336583 |
|  | LMNA | 15.40901632 | 1.248743278 |
|  | C3AR1 | 14.92290717 | 0.551025991 |
|  | TLR2 | 9.4232939 | 0.767708349 |
|  | RAP1A | 7.106411083 | 0.294887763 |
|  | ZFR | 3.105381571 | 0.310935775 |
|  | COL3A1 | 2.857957726 | 0.298002514 |
|  | COL1A1 | 2.333934487 | 0.1792467 |
|  | S100A8 | 2.04577801 | 0.113949073 |
|  | LSM5 | 1.63997989 | 0.225427155 |
|  | CFH | 1.583413177 | 0.195423902 |
|  | S100A9 | 1.354421064 | 0.392080053 |
|  | TLR4 | 1.335493175 | 0.366182584 |
|  | Fold change (CSF1 vs CSF10) |  | Standard deviation |
| SMA Type 2 Patient 18 | PDCD6 | 173.5143332 | 0.057659607 |
|  | TOLLIP | 156.1380931 | 0.431907451 |
|  | SMAD2 | 79.91170427 | 0.096224827 |
|  | ELP3 | 45.722541 | 0.381249406 |
|  | S100A9 | 41.78844081 | 0.095031657 |
|  | LMNB2 | 21.6469886 | 0.346593572 |
|  | GSTA1 | 18.78526285 | 0.595614823 |
|  | ZBTB10 | 12.21862039 | 0.762737035 |
|  | NDUFB11 | 3.298037236 | 0.021180594 |
|  | COL1A1 | 3.184279403 | 0.23708451 |
|  |  | Fold change (CSF 1 vs CSF13) | Standard deviation |
| SMA Type 2 Patient 2 | FCGR1A | 584.1383638 | 0.355577169 |
|  | ANXA5 | 531.8858819 | 0.147952986 |
|  | TFB1M | 258.0360206 | 0.095628785 |
|  | CLU | 225.2235497 | 0.551154577 |
|  | PHF3 | 221.150279 | 0.155725009 |
|  | SIRT1 | 190.8595581 | 0.200595582 |
|  | TMEM135 | 190.4471571 | 0.158214718 |
|  | CDKN1C | 189.0623708 | 0.250601367 |
|  | S100A9 | 135.0406671 | 0.006485973 |
|  | EP300 | 130.1434965 | 0.442934195 |
|  | SMAD2 | 118.502489 | 0.793600312 |
|  | ELAVL1 | 103.0557968 | 0.307191066 |
|  | TOLLIP | 102.7914085 | 0.549177745 |
|  | COL1A1 | 100.9872348 | 0.018083017 |
|  | ZNF25 | 96.99897278 | 0.242291511 |
|  | NDUFB11 | 92.66278432 | 0.856761758 |
|  | MRPL43 | 88.59719613 | 0.947594479 |
|  | LSM5 | 69.95014131 | 0.114391758 |
|  | LMNA | 53.77257768 | 0.108234293 |
|  | MBP | 46.42815795 | 0.040258552 |
|  | PDCD6 | 45.76825825 | 0.886061029 |
|  | ZBTB10 | 21.14065574 | 0.413692515 |
|  |  | Fold change (CSF1 vs CSF11) | Standard deviation |
| SMA Type 2 Patient 10 | FCGBP | 417.6680263 | 0.010656394 |
|  | PHF3 | 233.3443559 | 0.579730693 |
|  | CDKN1C | 204.7861992 | 0.244733551 |
|  | SIRT3 | 58.93410061 | 0.333015907 |
|  | SMAD2 | 47.80902195 | 0.486909747 |
|  | MBP | 22.49584673 | 0.019454875 |
|  | TERF2 | 11.02807011 | 0.094799276 |

***Table 2. Gene lists of upregulated and downregulated genes from qPCR array***
|  | SIRT6 | 4.794880595 | 0.191212338 |
| --- | --- | --- | --- |
|  | Fold change (CSF1 vs CSF9) |  | Standard deviation |
| SMA Type 3 Patient 7 | S100A8 | 98.42173392 | 0.043989086 |
|  | POLRMT | 80.29445387 | 0.475298452 |
|  | SIRT1 | 77.8899168 | 0.146657723 |
|  | TINF2 | 70.94753706 | 0.015234193 |
|  | SIRT6 | 46.80597869 | 0.670577912 |
|  | FCER1G | 40.38169017 | 0.416247735 |
|  | LMNB2 | 37.95351574 | 0.333048049 |
|  | SMAD2 | 34.6404783 | 0.693343932 |
|  | RAP1A | 31.79333196 | 0.962911765 |
|  | CDKN1C | 7.696561994 | 0.210202678 |
|  | COL3A1 | 1.970087049 | 0.034445243 |
| Downregulated genes from qPCR array |  |  |  |
| Patient | Gene | CSF samples |  |
|  |  | Fold change (CSF1 vs CSF8) | Standard deviation |
| SMA Type 1 Patient 1 | ANGEL2 | 0.47560534 | 0.324373876 |
|  | C1QB | 0.217218762 | 2.147239377 |
|  | ELAVL1 | 0.016014869 | 1.494078081 |
|  | SIRT1 | 0.010019592 | 0.149775637 |
|  | ZBTB10 | 0.009851439 | 0.149775637 |
|  | WRN | 0.007996542 | 0.149775637 |
|  | SIRT3 | 0.007923551 | 0.149775637 |
|  | MBP | 0.005150489 | 0.06821062 |
|  | Fold change (CSF1 vs CSF4) |  | Standard deviation |
| SMA Type 1 Patient 6 | COL3A1 | 0.644604761 | 0.240358893 |
|  | C1QB | 0.582525045 | 0.452585549 |
|  | SNAP23 | 0.378493734 | 0.309616138 |
|  | MBP | 0.3273256 | 0.097744264 |
|  | ANGEL2 | 0.306683707 | 0.698132674 |
|  | LMNA | 0.205893418 | 0.249621848 |
|  | JAKMIP3 | 0.156796644 | 0.365855905 |
|  | TXNIP | 0.063395626 | 0.15679019 |
|  | C5AR1 | 0.054662431 | 0.15679019 |
|  | BUB1B | 0.050576119 | 0.15679019 |
|  | ZBTB10 | 0.020336097 | 0.420825512 |
|  | EML1 | 0.019549467 | 0.15679019 |
|  | ZFR | 0.01164457 | 0.15679019 |
|  | TINF2 | 0.009555146 | 0.15679019 |
|  | ELAVL1 | 0.00902345 | 0.15679019 |
|  | SIRT3 | 0.006730763 | 0.15679019 |
|  | Fold change (CSF1 vs CSF5) |  | Standard deviation |
| SMA Type 1 Patient 17 | C1QB | 0.421290775 | 1.849573585 |
|  | ANGEL2 | 0.298832887 | 0.040388811 |
|  | FCGR1A | 0.283775686 | 0.285136655 |
|  | FCER1G | 0.152380734 | 0.092837075 |
|  | NDUFB11 | 0.142123082 | 0.33653961 |
|  | JAKMIP3 | 0.020542569 | 2.049182279 |
|  | LYZ | 0.019036281 | 0.116481975 |
|  | PDCD6 | 0.009608537 | 0.116481975 |
|  | ZNF25 | 0.008402913 | 0.116481975 |
|  | WRN | 0.007055008 | 0.116481975 |
|  | TFB1M | 0.005031118 | 0.116481975 |
|  | Fold change (CSF1 vs CSF10) |  | Standard deviation |
| SMA Type 2 Patient 18 | ANGEL2 | 0.265039053 | 0.103281794 |
|  | CLU | 0.16171658 | 0.149634744 |
|  | WRN | 0.061251482 | 0.038688432 |
|  | CD163 | 0.038358619 | 0.038688432 |
|  | FCGR1A | 0.019700529 | 0.038688432 |
|  | MRPL43 | 0.016099484 | 1.606029444 |
|  | ARL6IP6 | 0.011497917 | 0.038688432 |
|  | RAP1A | 0.009820707 | 0.038688432 |
|  | Fold change (CSF 1 vs CSF13) |  | Standard deviation |
| SMA Type 2 Patient 2 | ANGEL2 | 0.574657434 | 0.150396102 |
|  | TINF2 | 0.218425785 | 1.196952087 |
|  | CD14 | 0.116463332 | 1.594650575 |
|  | ARID1A | 0.014563725 | 0.012236546 |
|  | CD163 | 0.013182965 | 0.012236546 |
|  | VWA5A | 0.006281219 | 0.012236546 |
|  |  | Fold change (CSF1 vs CSF11) | Standard deviation |
| SMA Type 2 Patient 10 | C1QB | 0.915402274 | 0.114177078 |
|  | ANGEL2 | 0.498470727 | 0.042179981 |
|  | C1QC | 0.403954375 | 1.127620478 |
|  | S100A8 | 0.121075818 | 0.015522779 |
|  | COL3A1 | 0.077984591 | 0.345294375 |
|  | FCGR1A | 0.05878036 | 0.104202059 |
|  | C1QA | 0.055116066 | 0.104202059 |
|  | HSF1 | 0.035261923 | 0.104202059 |
|  | RAP1A | 0.02382736 | 0.104202059 |
|  | CASP1 | 0.022797964 | 0.104202059 |
|  | VWA5A | 0.012304053 | 0.104202059 |
|  | WRN | 0.011627615 | 0.104202059 |
|  | ANXA5 | 0.008541818 | 0.104202059 |
|  | TMEM135 | 0.008306068 | 0.104202059 |
|  | COL1A1 | 0.006265169 | 0.256562015 |
|  | S100A9 | 0.002258564 | 0.104202059 |
|  | Fold change (CSF1 vs CSF9) |  | Standard deviation |
| SMA Type 3 Patient 7 | ANGEL2 | 0.333549289 | 0.215145315 |
|  | FCGBP | 0.216368666 | 1.906835713 |
|  | ZFR | 0.106991425 | 1.087050383 |
|  | TERF2 | 0.09783228 | 1.128964282 |
|  | TLR4 | 0.059815743 | 1.582313679 |
|  | SIRT3 | 0.030033526 | 0.037515975 |
|  | COL1A1 | 0.023115625 | 1.196391903 |
|  | PHF3 | 0.016128083 | 0.037515975 |
|  | MRPL43 | 0.014311411 | 0.037515975 |
|  | ZNF25 | 0.009322887 | 0.037515975 |
|  | FCGR1A | 0.00811128 | 0.037515975 |
|  | PDCD6 | 0.006029665 | 0.037515975 |
|  | ANXA5 | 0.005820223 | 0.037515975 |

**Figure 1.**
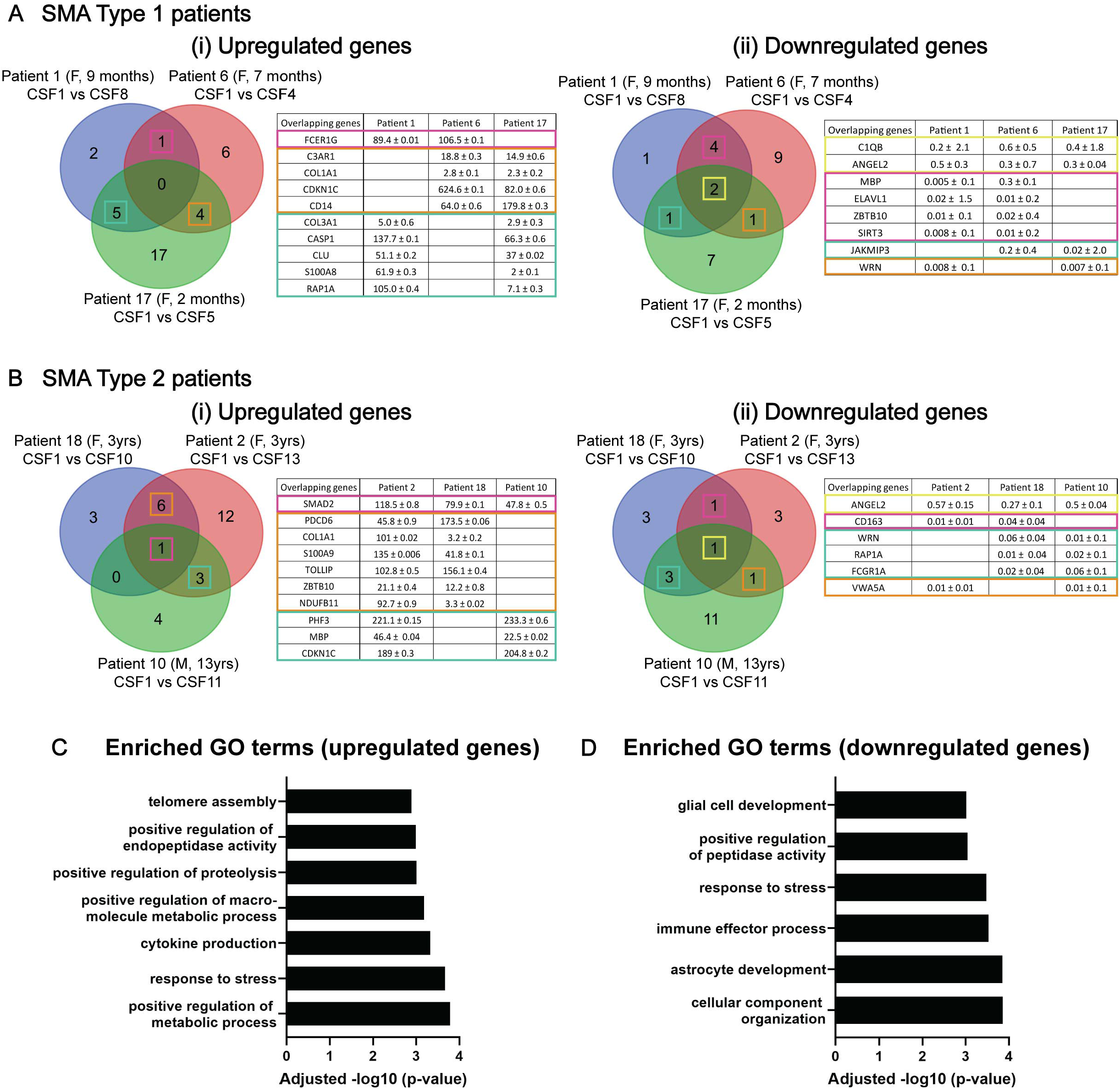
Screening for mRNA biomarkers in SMA patient CSF samples using a RT-qPCR array. Pre-amplified cDNA samples from SMA Type 1 **(A)** and Type 2 **(B)** patient CSF samples were run on RT^2^ Profiler PCR array (SMA Type 3 patient data included in Table 2). Upregulated **(i)** and downregulated **(ii)** genes were detected; gene expression of latest post-nusinersen treated CSF samples (e.g. CSF 11) was normalized to pre-treatment gene expression (i.e. CSF1) for each patient. The Venn diagrams show the number of overlapping genes and genes that are specific to each patient, and the tables detail the overlapping genes and their relative fold changes ± standard deviation (SD) compared to CSF1 **(A-B)**. Gene ontology analyses were performed on upregulated genes **(C)** and downregulated genes **(D)** across all SMA patient samples.

We next assessed microRNAs as potential biomarkers in the SMA pre- and post-nusinersen treated CSF samples, focusing on those that have been previously been implicated in SMA disease pathology (Fig. 2). We observed an overall upregulated trend in expression across all microRNAs in patient samples, with CSF1 represents the baseline microRNA expression prior to treatment (dotted lines in each graph). In some patients there was an earlier increase in microRNA expression (detected in CSF4/5 samples), including miR-132 (Fig. 2A), miR-218 (Fig. 2B) and miR-9 (Fig. 2C), whereas other microRNAs showed a more gradual change in expression across the nusinersen treated samples (Fig. 2D-F). This trend was mainly observed in Type 1 and 2 patient samples, who clinically responded well to treatment (Table 1). This is an encouraging trend for microRNAs that are typically downregulated in SMA, including miR-132, miR-218, miR-9 and miR-23a (Fig. 2A-D). miR-431 expression is increased in SMA pathology, but it appears to remain relatively unchanged post-nusinersen treatment due to the low fold change levels in expression observed (Fig. 2E). Similarly, increased miR-146a expression is associated with SMA pathology; however, post-nusinersen treated CSF samples in SMA patients continued to show increased expression of this microRNA (Fig. 2F).

**Figure 2.**
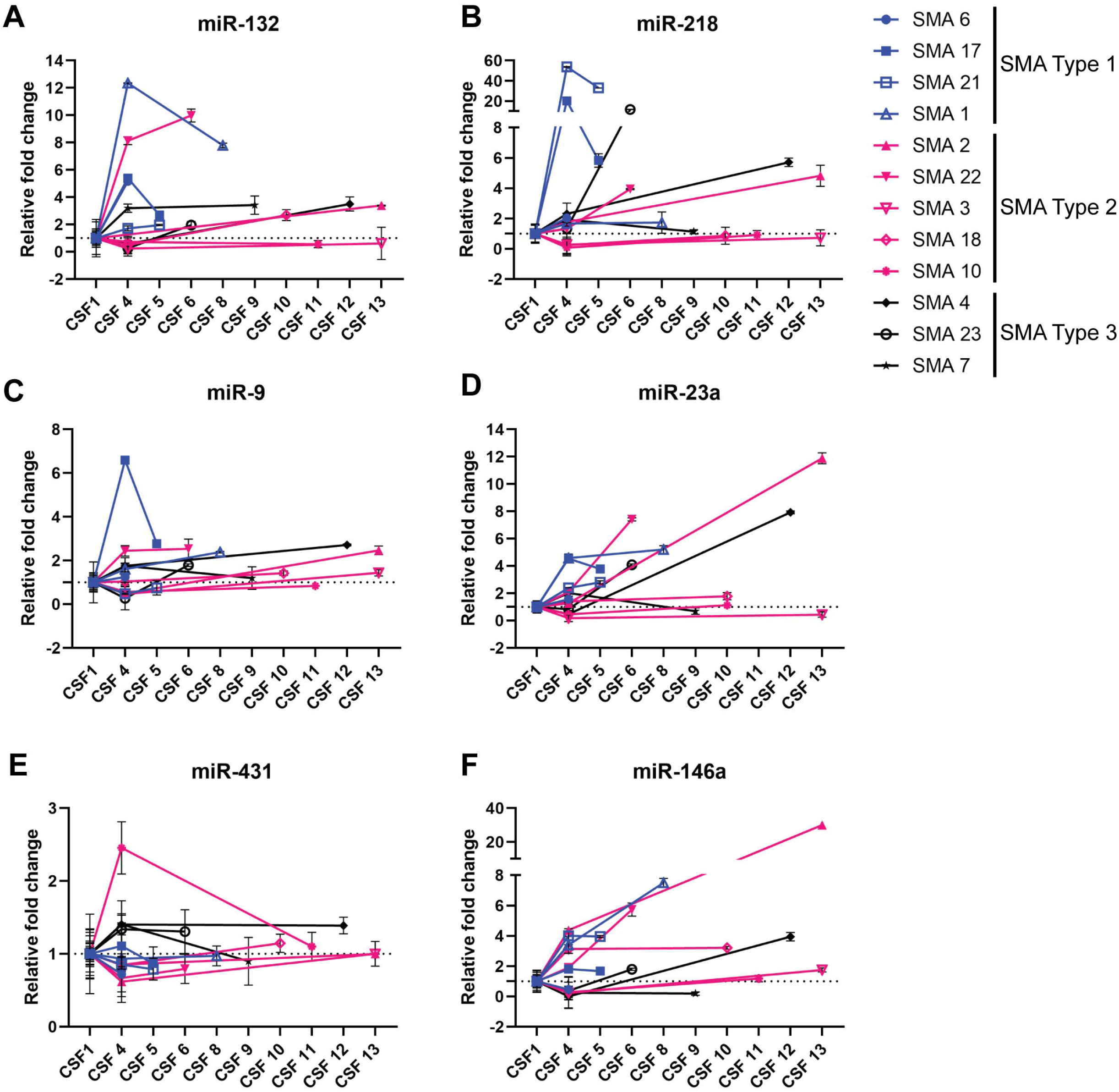
Screening for microRNA biomarkers in SMA patient CSF samples via RT-qPCR. Pre-amplified cDNA samples from SMA Type 1 **(blue data points)**, Type 2 **(pink data points)** and Type 3 **(black data points)** patient CSF samples were screened for previous microRNAs implicated in SMA disease pathology. miR-132 **(A)**, miR-218 **(B)**, miR-9 **(C)**, miR-23a **(D)**, miR-431 **(E)** and miR-146a **(F)** expression is represented in the graphs as relative fold change; CSF4 and latest post-nusinersen treated CSF samples were normalized to pre-treatment samples (CSF1).

We have previously found that astrocyte-produced miR146a induces a reduction in choline acetyltransferase (ChAT) expression and neurite outgrowth in iPSC-derived motor neuron cultures (19). Due to the detection of inflammatory and astrocyte and glial development GO terms and the increased expression of astrocyte-associated miR-146a in post-nusinersen treated CSF samples, we used mRNA sequencing to further dissect molecular mechanisms that may be contributing to motor neuron malfunction and loss in SMA. Principal component analysis of all motor neuron samples demonstrates clustering by differentiation culture replicates (Fig. 3A). Importantly, miR146a treated samples in all clusters closely align with untreated (UTX) or negative mimic treated samples, which was also reflected in the DESeq2 comparative analysis where no statistically significant differentially expressed genes were determined (-log10 adjusted p value < 0.05). Previous reports have suggested that microRNAs are responsible for fine-tuning gene expression and thus microRNA expression modification may have more subtle effects (24). We therefore analyzed genes without the p value correction (p value < 0.05), but with robust fold change in expression (log2FC <-1 or >1) to identify 163 and 121 differentially expressed genes comparing healthy UTX vs miR-146a treated samples and SMA UTX vs SMA mir146a treated samples, respectively (Table 3). In both datasets, a larger number of downregulated genes were observed (149 genes healthy UTX vs healthy miR-146a treated; 115 genes SMA UTX vs SMA mir146a treated). GO analyses on these downregulated genes showed enrichment of terms related to extracellular matrix (ECM), metallopeptidase activity, collagen-containing ECM, catabolic processes and plasma membrane components (Fig. 2B and C), with heatmaps highlighting downregulated genes found in the GO terms (Fig. 2D). Of particular interest was the identification of genes encoding matrix metalloproteinases (MMP17, 10, 1), a disintegrin and metalloproteinases (ADAM29) with thrombospondin repeats (ADAMTS2, 9, 3 14, ADAMTSL2), and other ECM-related genes (e.g. TNC, THBS2, HAPLN1, NEDD9, ITGA4) because of their known role at the perineuronal net and interactions at the synapse. This led us to look further for synaptic related genes within the downregulated datasets, to which we identified genes associated with dendritic spine formation and stability (SYT6, GPX3), axonal guidance and outgrowth (PLXNB3, NGEF,SPON2), glutamate transporters and receptor subunits (SLC17A8, GRID2, GRM8, GRIN3A) and potassium voltage gated ion channels (KCNE4, KCNQ3, KCNQ5). We also found a downregulation of genes needed to maintain neuronal health against oxidative damage (GPX3, ENC1) and through neurotrophic factor signaling (NTRK1, GFRA2). Intriguingly, synapse was an enriched GO term when analyzing known miR-146a gene targets (Fig. 2F); among these established miR-146a targets, we identified the vesicular glutamate transporter, SLC17A8, within our downregulated gene sets and identified genes with similar functions to known miR-146a targets (e.g. MMP and ADAMTS proteases, and SYT6).

**Table 3.**
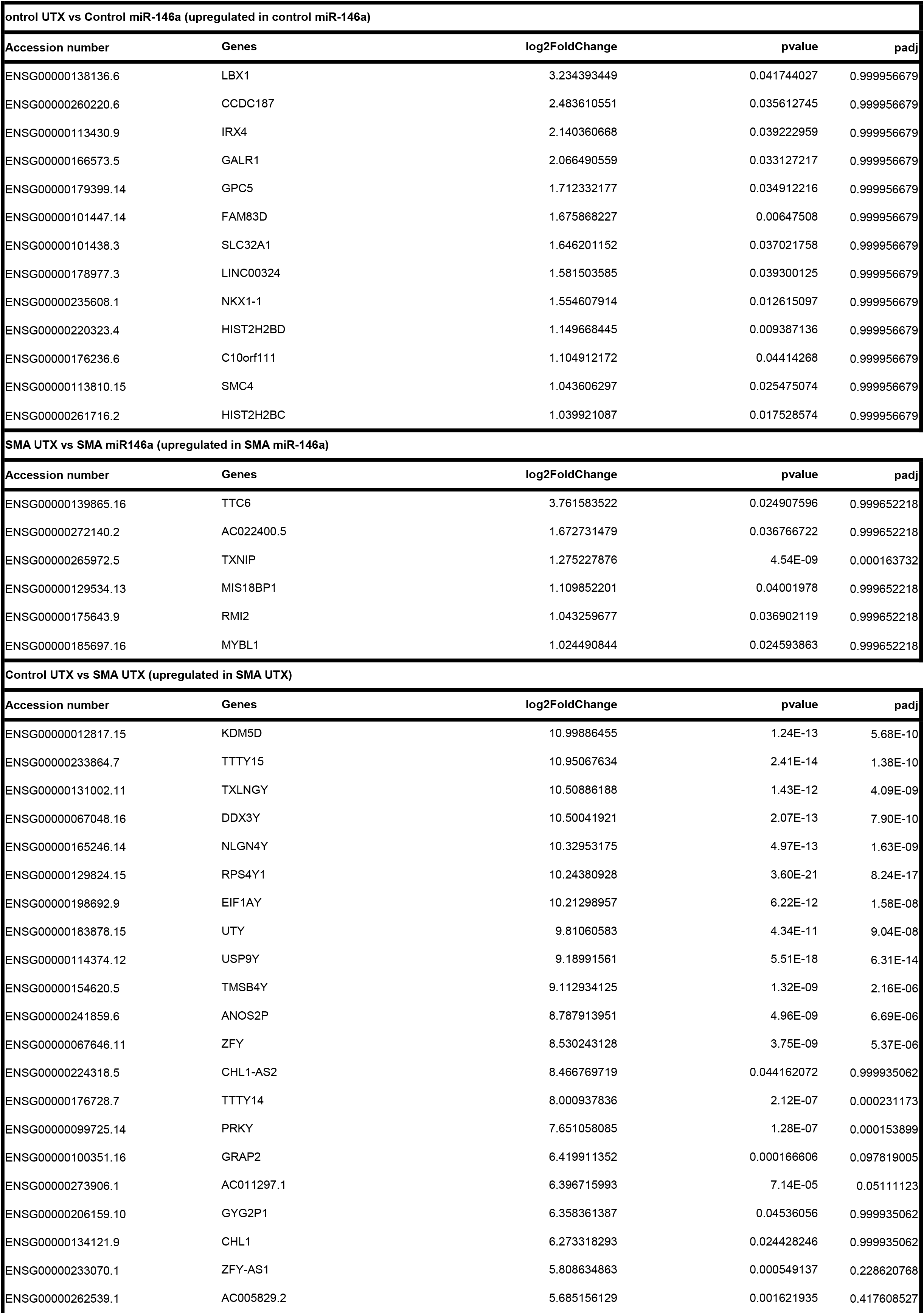
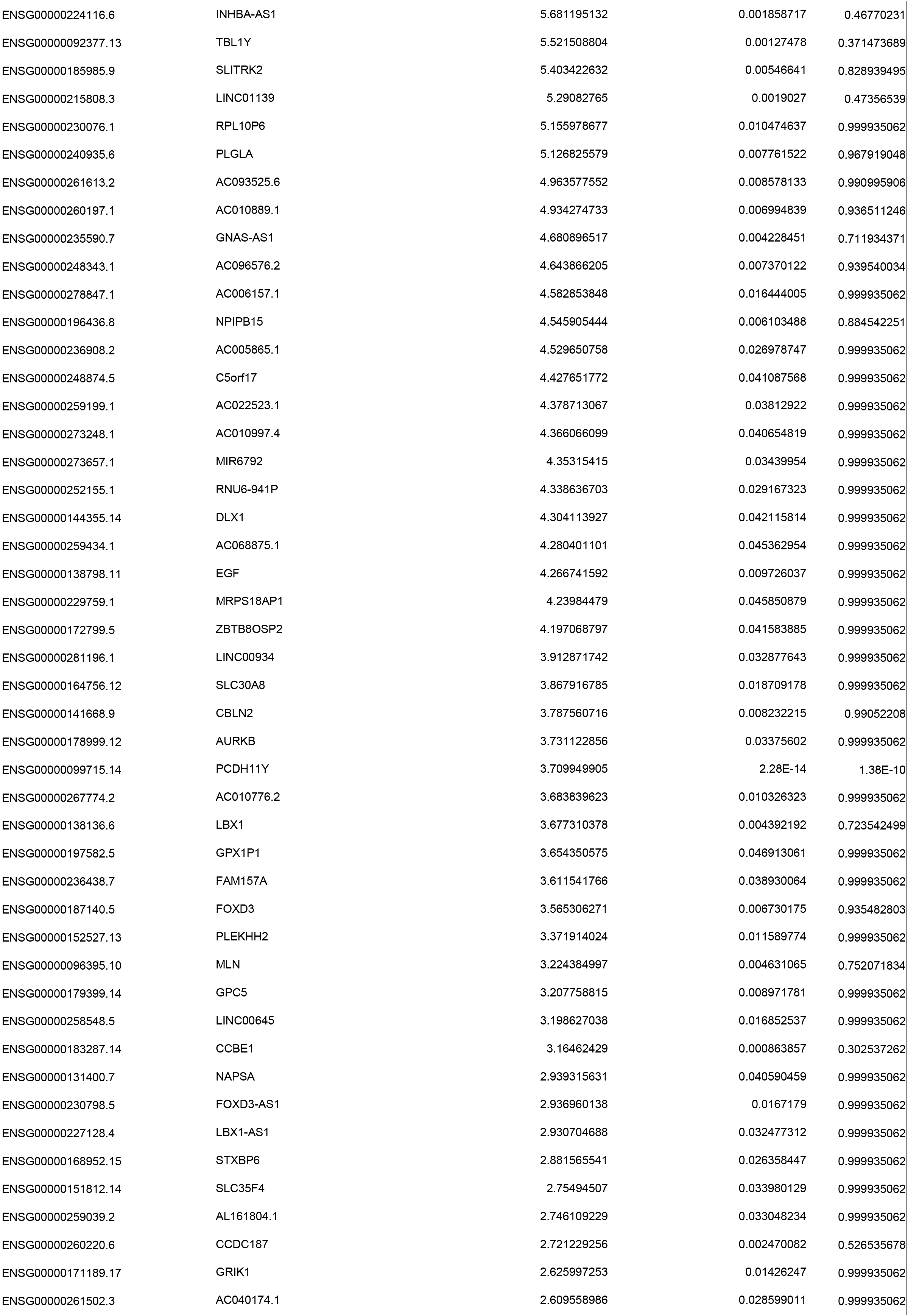
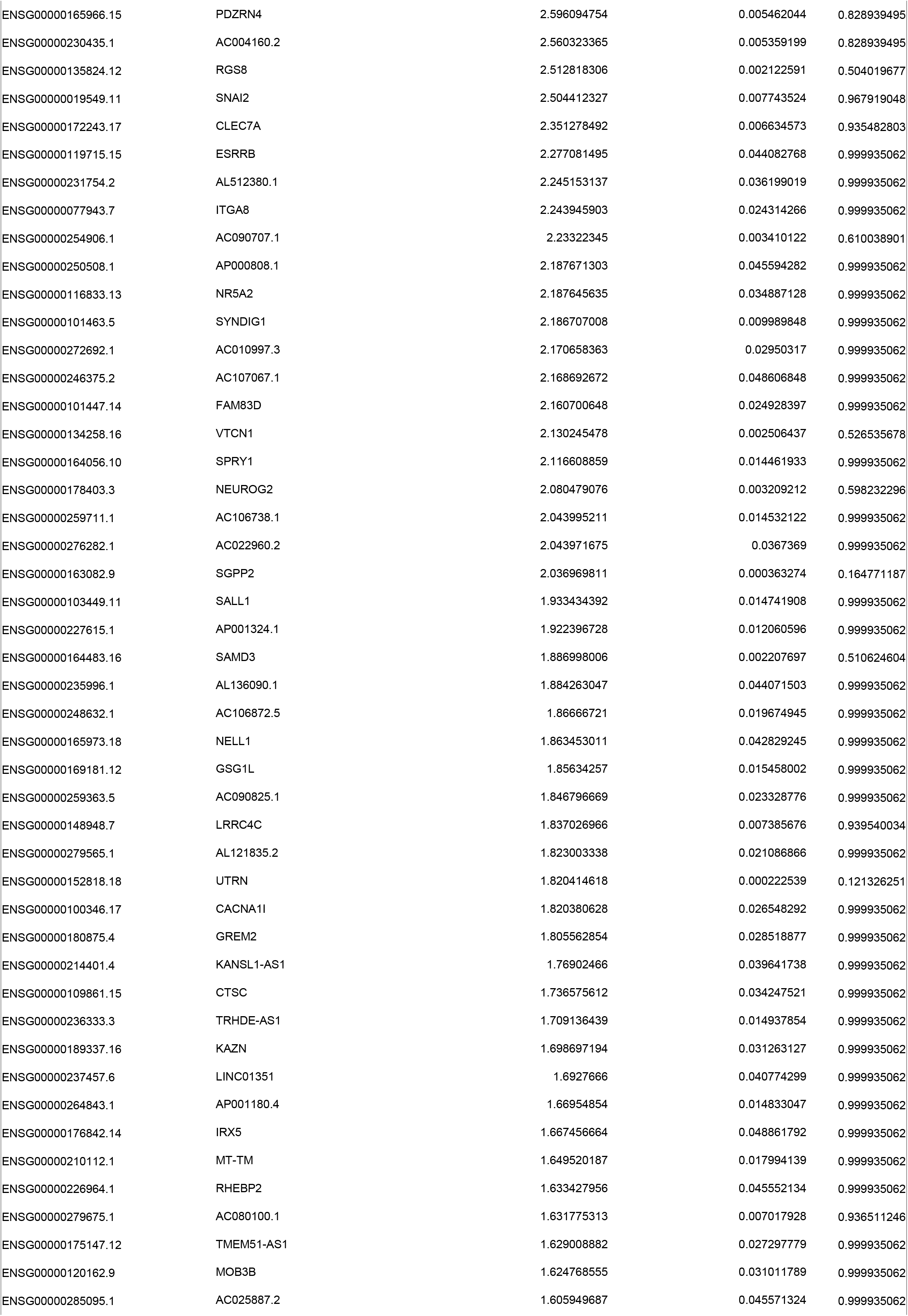
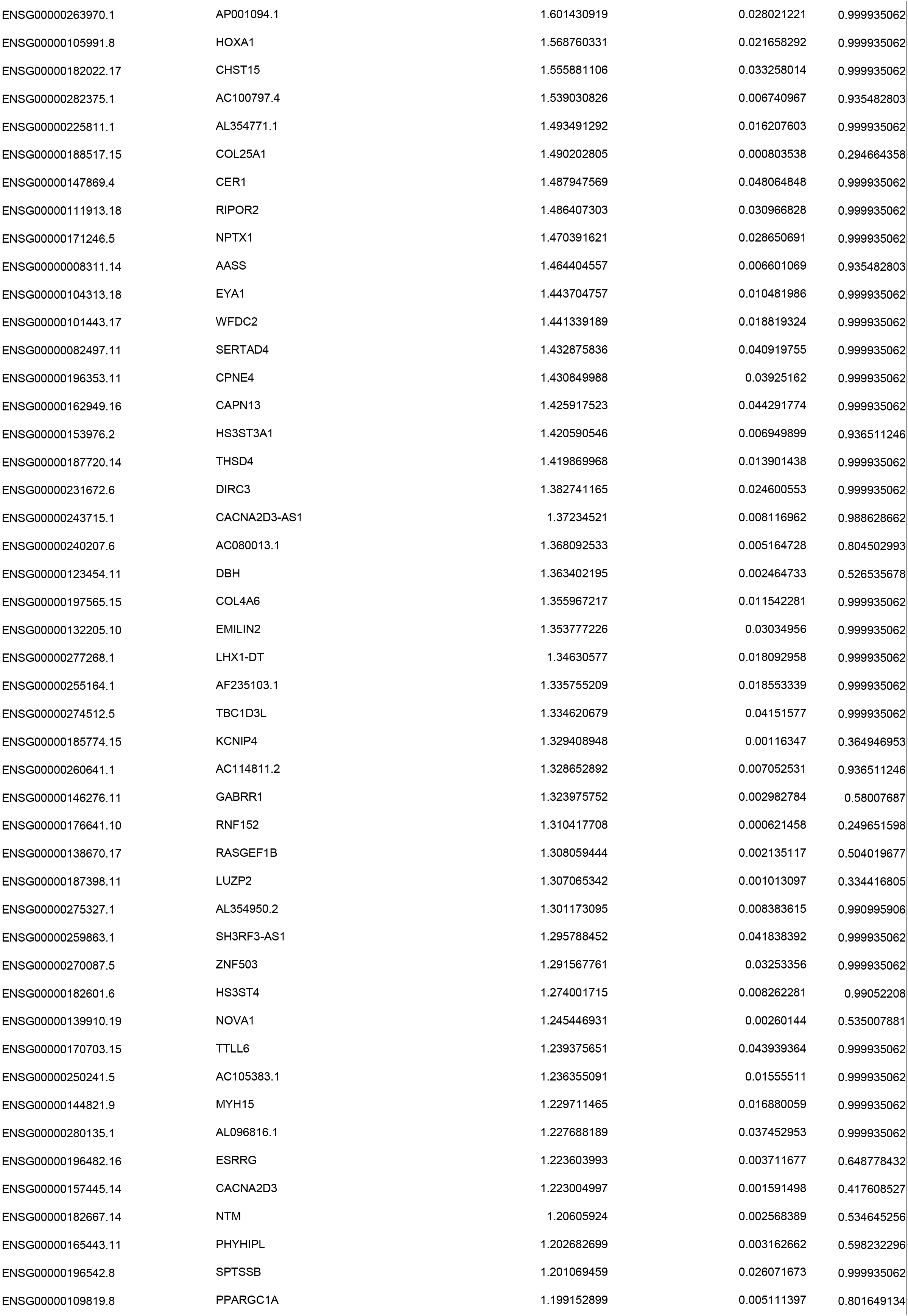
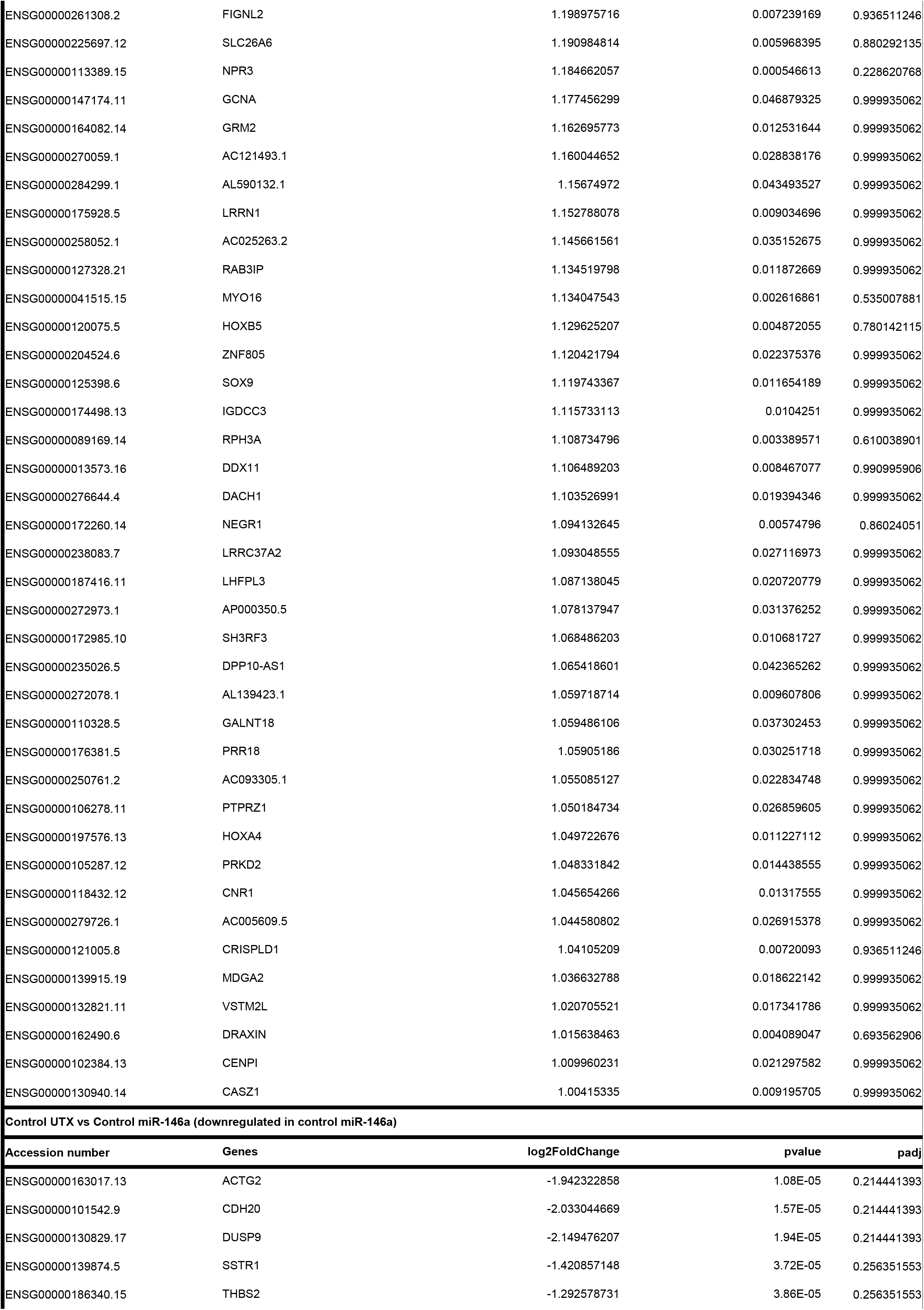
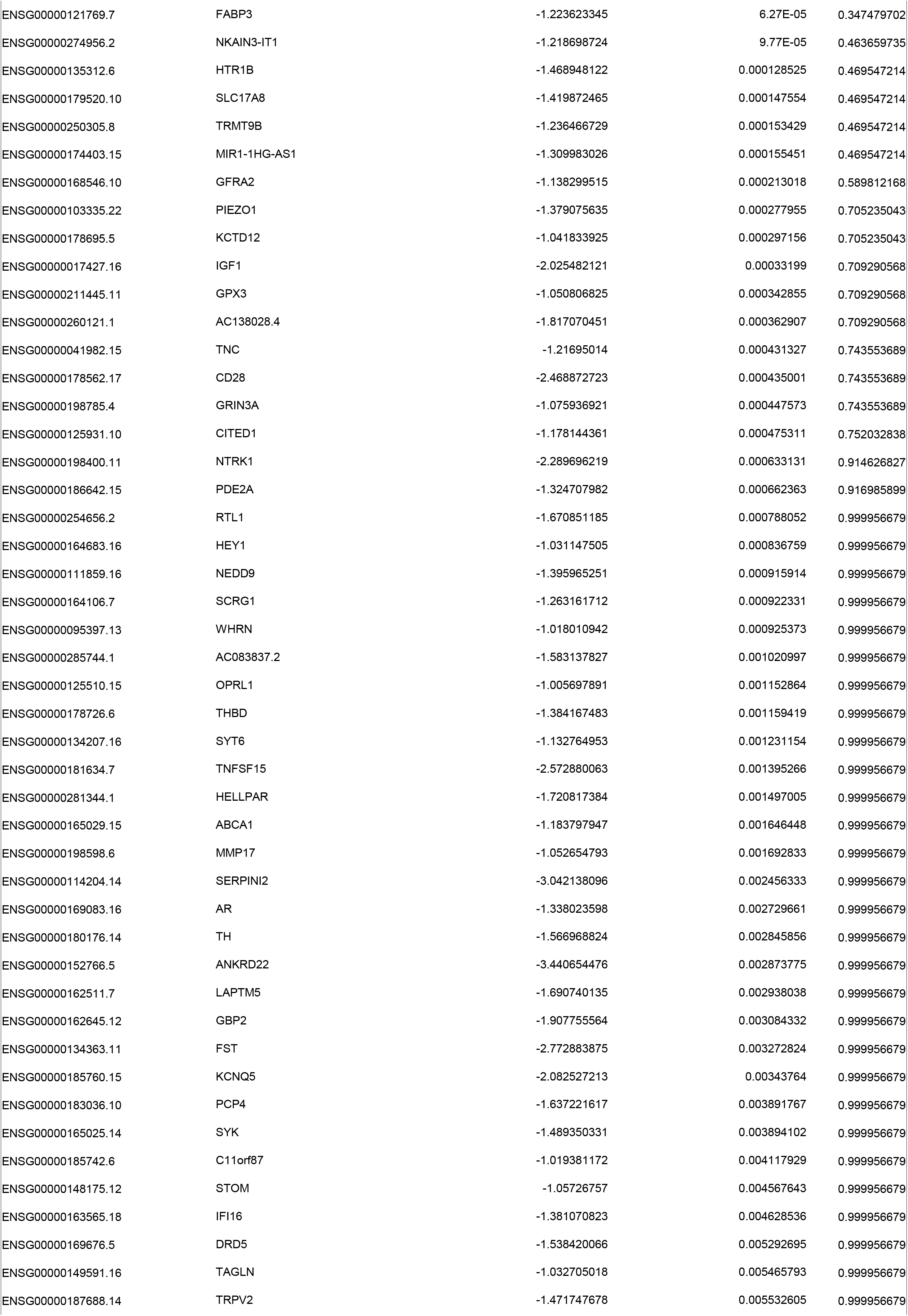
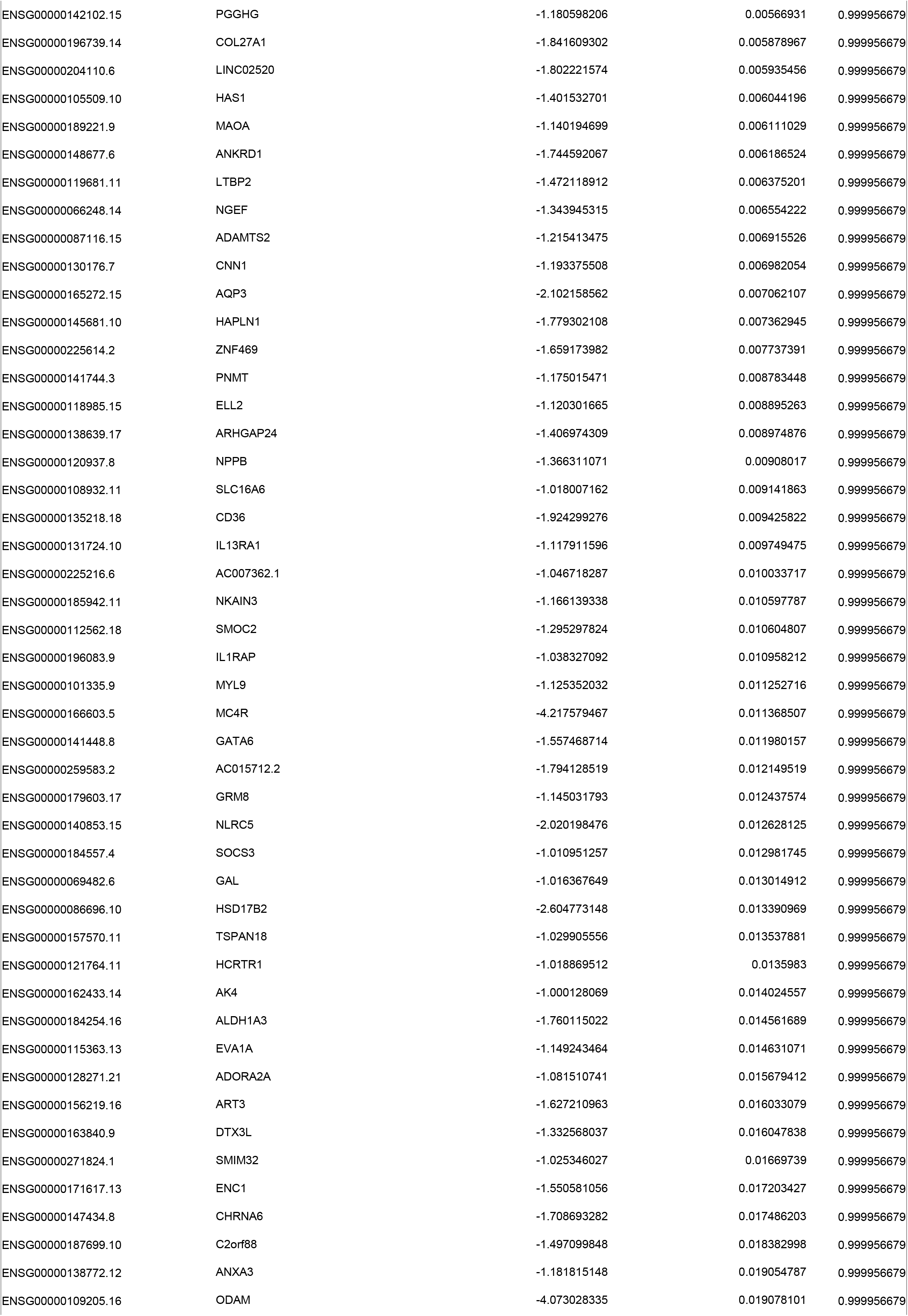
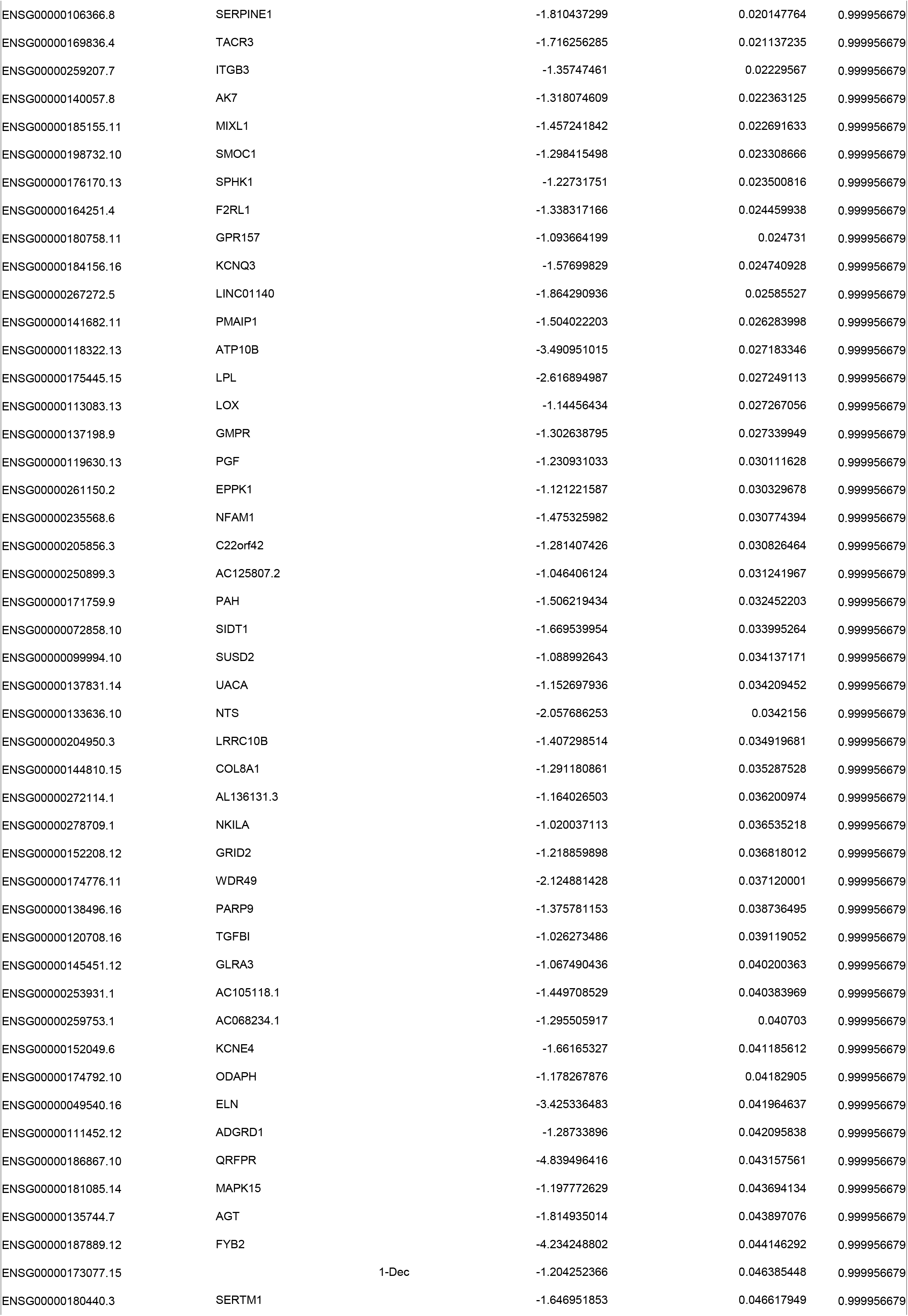
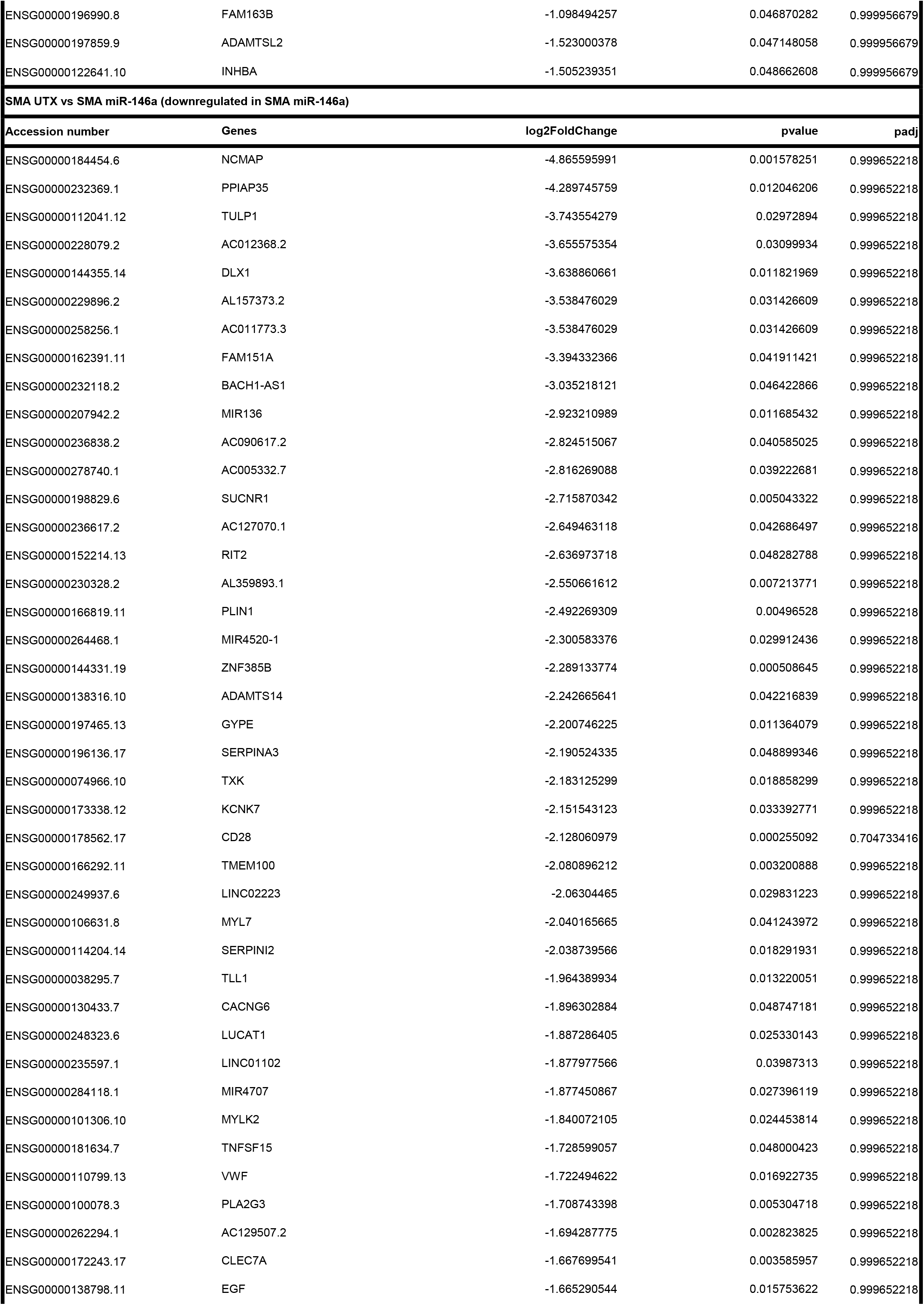
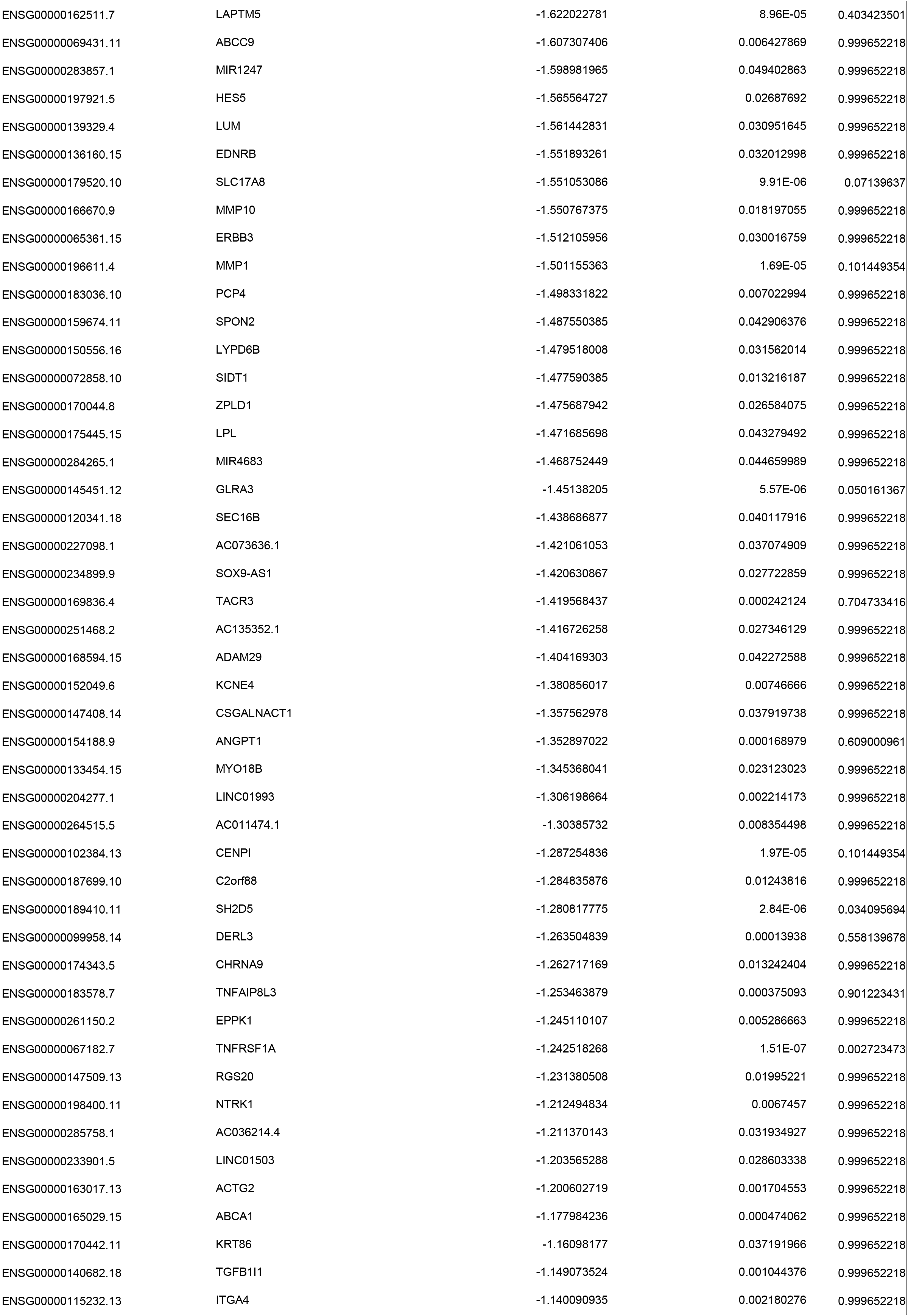
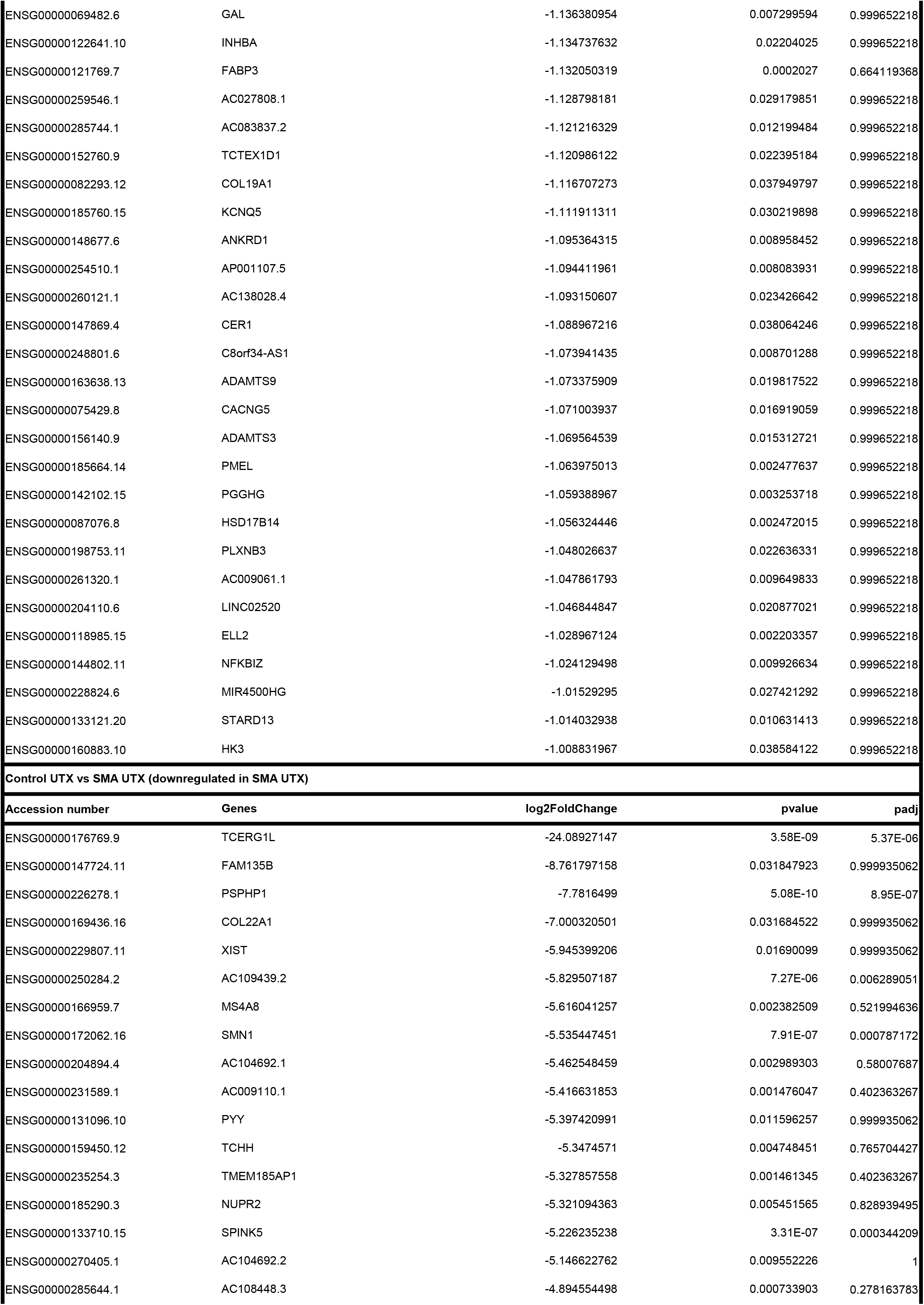
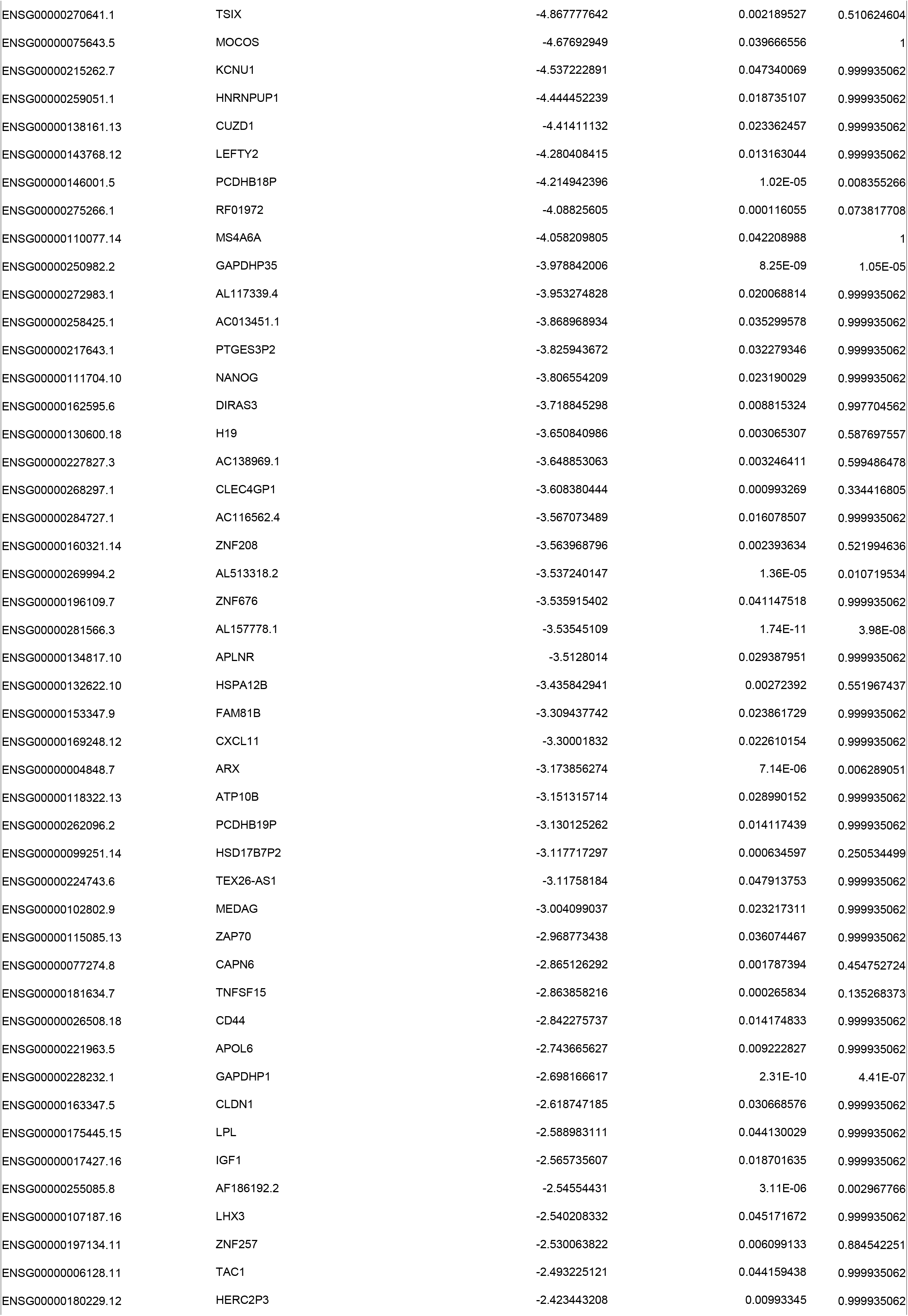
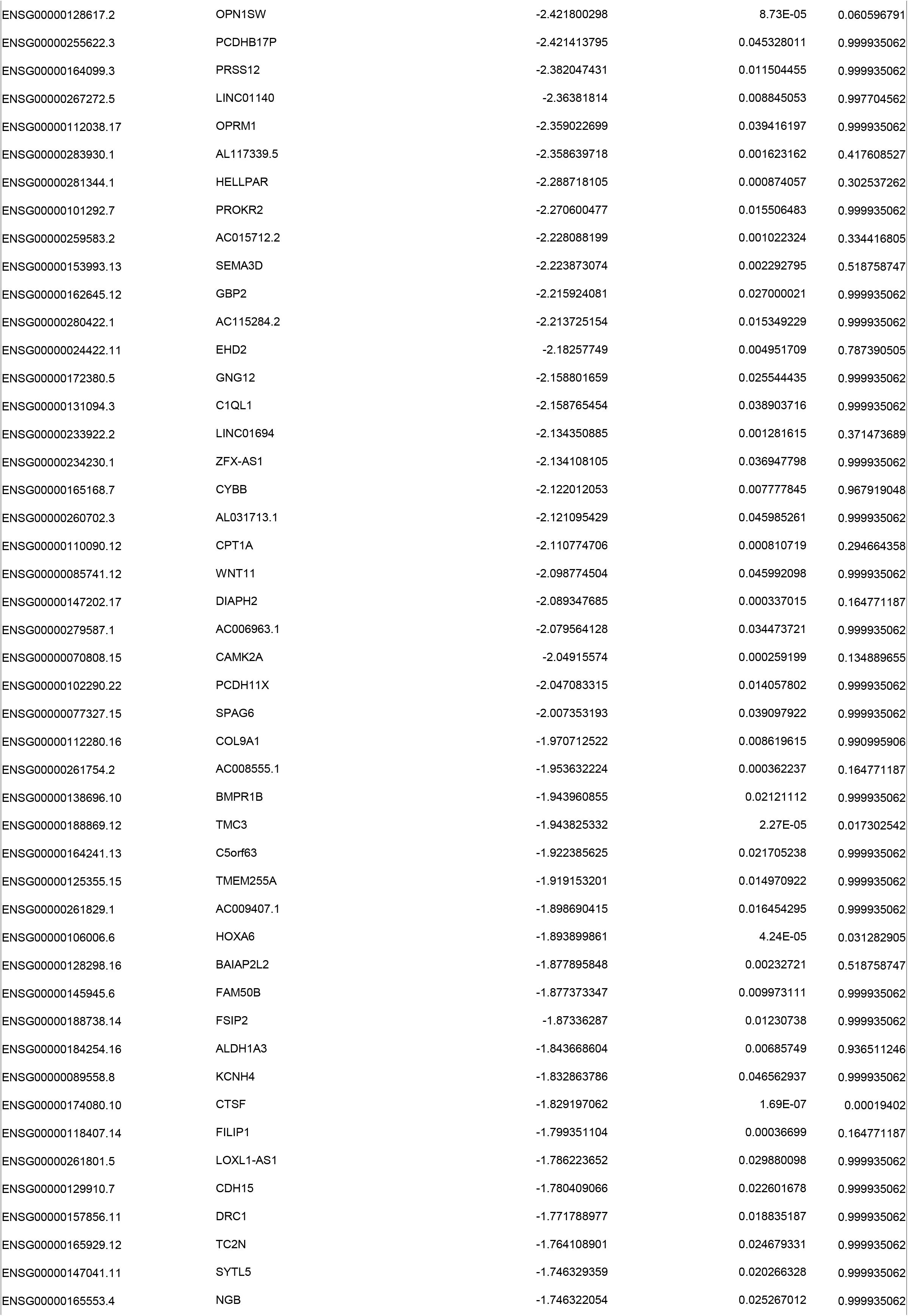
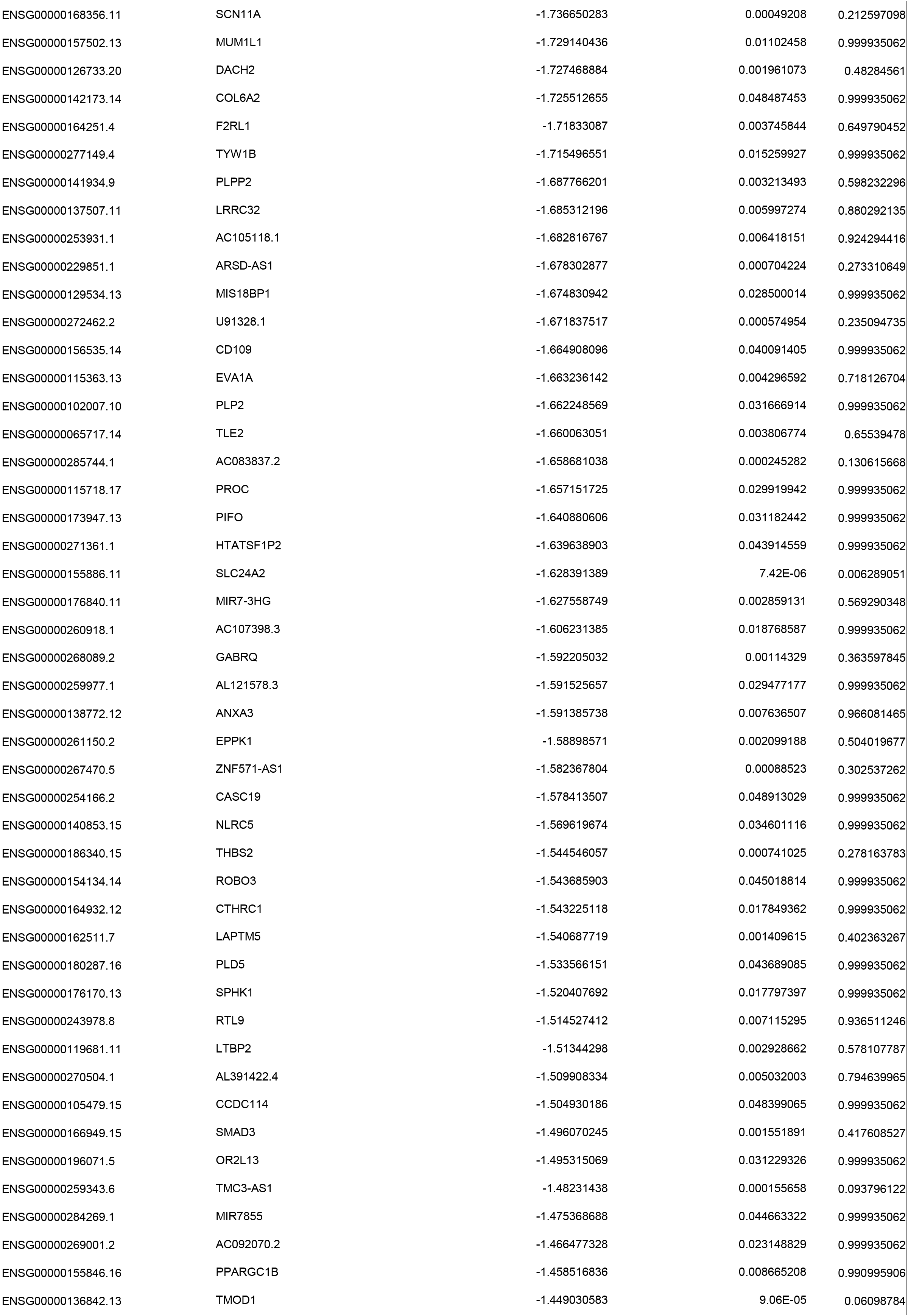
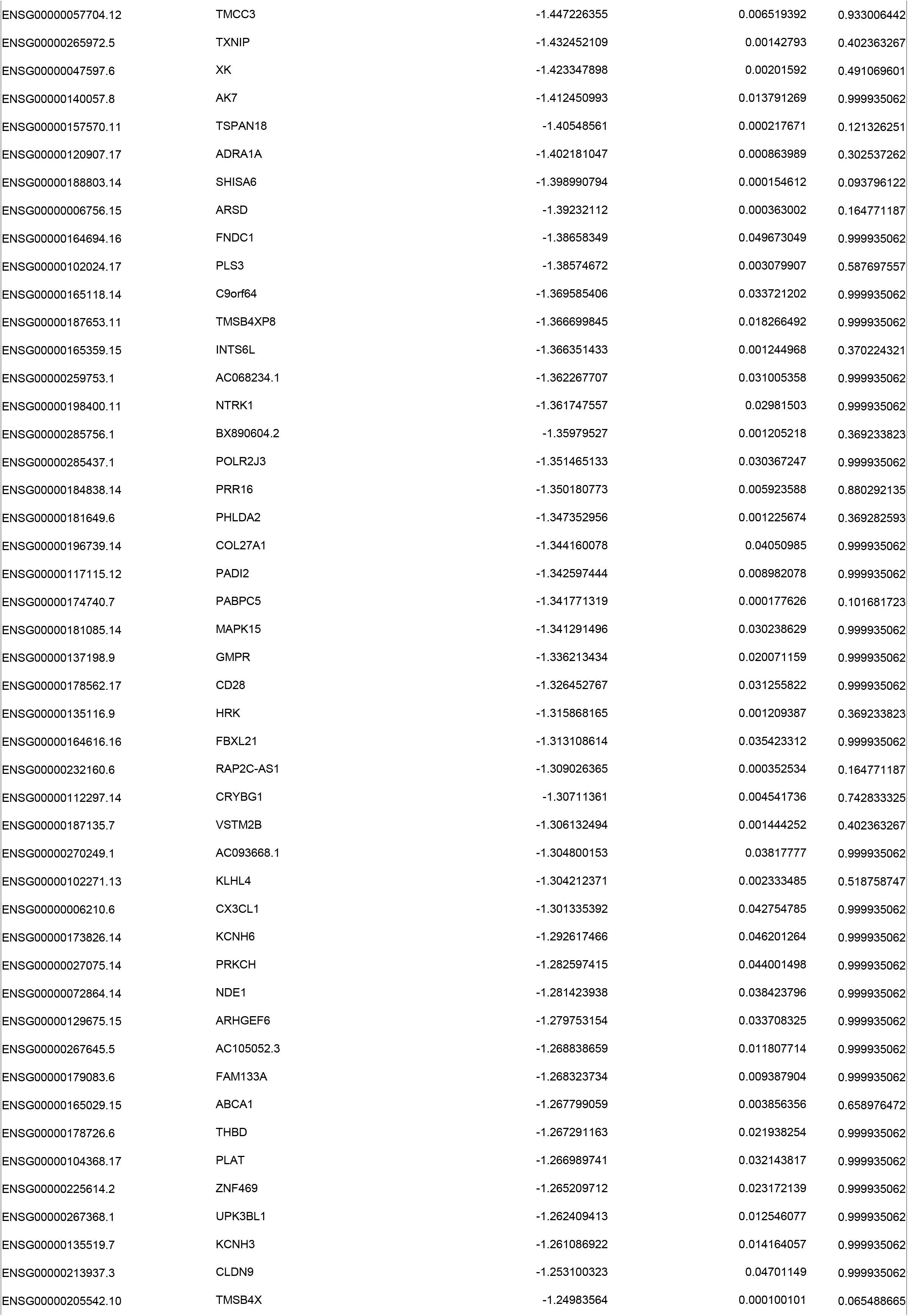
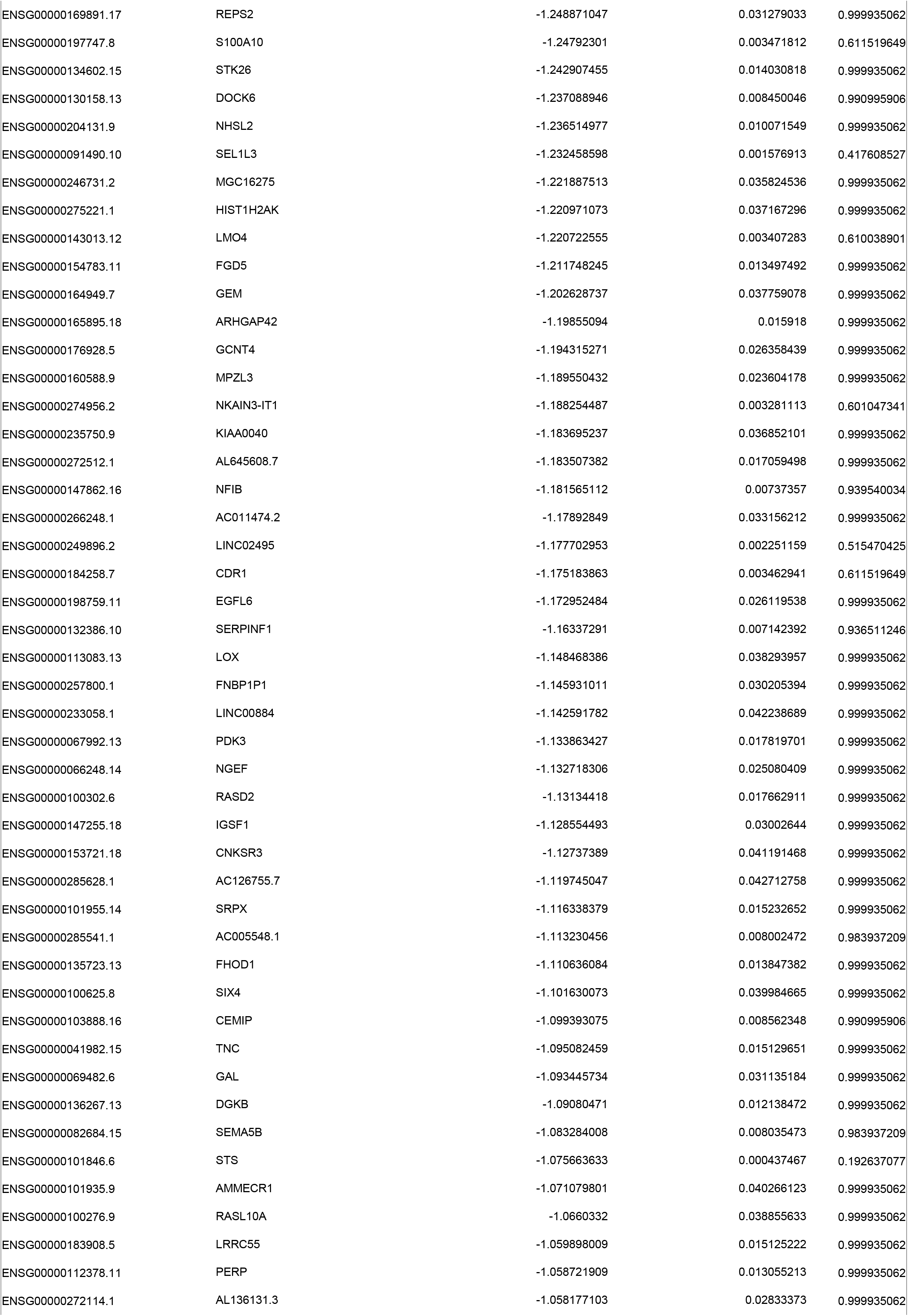
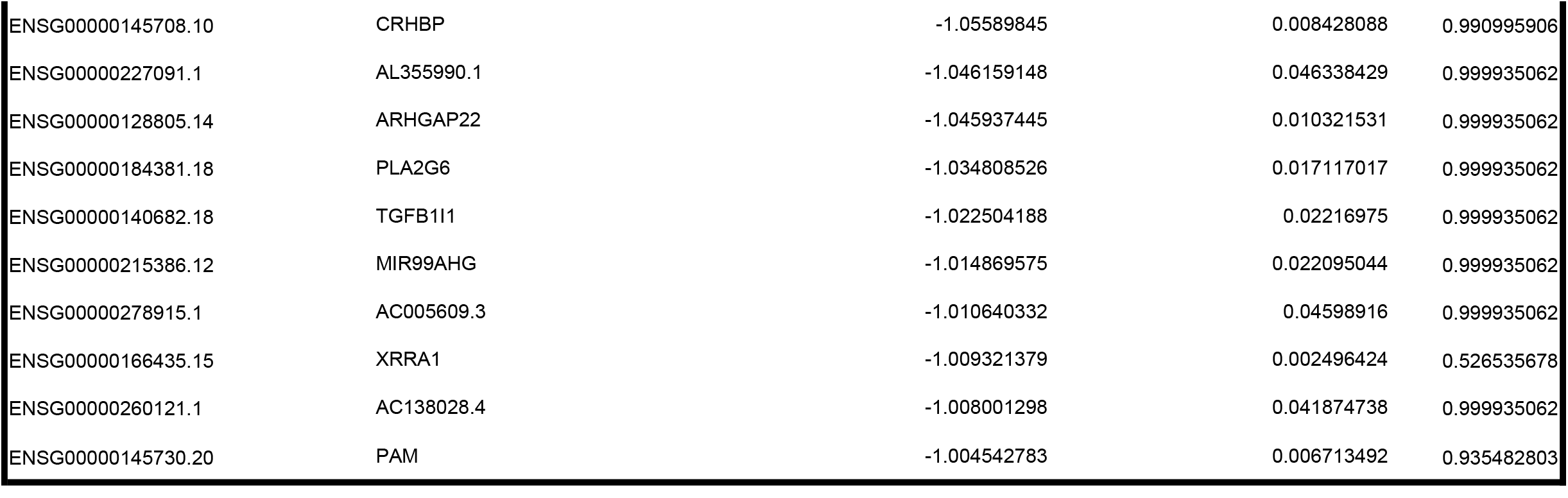
RNA seq differentially expressed genes (healthy UTX vs healthy miR-146a, SMA UTX vs SMA miR-146a, healthy UTX vs SMA UTX). padj; adjusted p-value.

**Figure 3.**
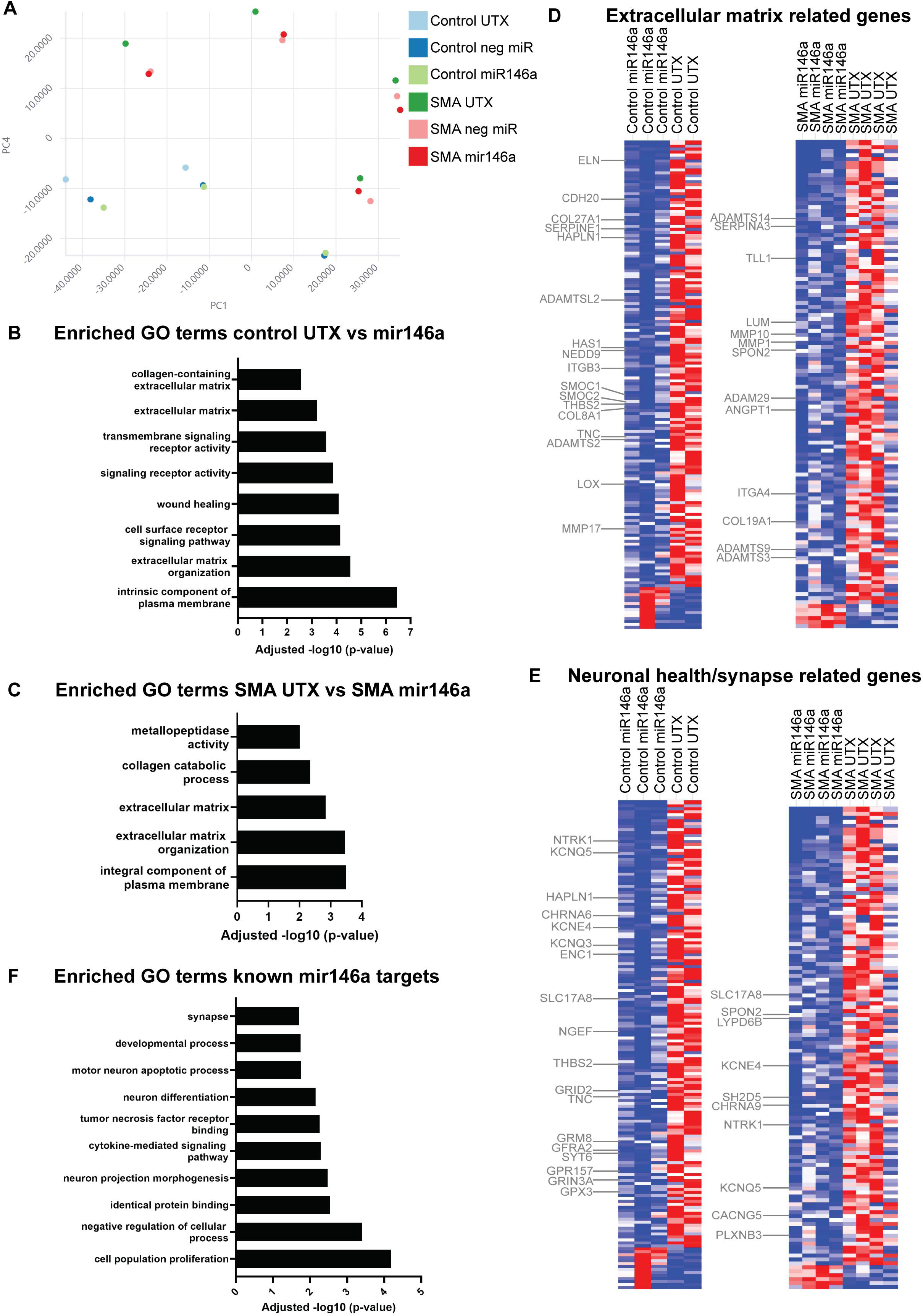
RNA seq analysis of iPSC-derived motor neurons with mir146a treatment. DeSeq2 analysis was performed on all motor neuron samples including untreated (UTX), negative miR mimic (neg miR) and miR-146a treated from both control and SMA iPSC lines. This incorporates a Principal Component Analysis **(A)** to assess the variance between samples based on gene expression. Gene ontology (GO) analysis was performed on those differentially expressed genes (p value<0.05) that were downregulated (log2 fold change) in control UTX vs miR-146a treated samples **(B)** and SMA UTX vs SMA miR-146a treated samples **(C)**. The heatmaps highlight those genes encoding extracellular matrix proteins (some related to the synapse) **(D)** and genes encoding neuronal health/synapse related proteins **(E)** that are downregulated in control and SMA miR-146a treated samples, respectively. GO analysis was additionally performed on a gene list of known miR-146a targets **(F)** which also showed an enrichment of GO terms associated with synapse and neuron projection morphogenesis.

To assess if the downregulation of synaptic genes due miR-146a exposure could impact motor neuron function, we used a multi-electrode array (MEA) approach to assess spontaneous action potentials in UTX, negative mimic and miR146a mimic treated iPSC-derived motor neurons (Fig. 4A). Motor neuron action potentials were detected beginning at 8 days after plating (Fig. 4B). Prior to treatment, there was no significant difference observed in the weighed mean firing rate (WMFR) between healthy and SMA motor neurons, although we did observe a slight increase in the SMA motor neuron activity (Fig. 4C). After 48 hours of miR-146a treatment, WMFR decreased in the healthy motor neurons whereas SMA motor neurons showed increased activity (Fig. 4D). Although not statistically significant, these data still represent an interesting observation regarding intrinsic synaptic differences between healthy and SMA motor neurons. We returned to the transcriptome data to compare healthy vs SMA motor neuron samples to look for difference in synaptic gene expression (Fig 4E; Table 3). Similar to the miR-146a treated samples, we saw a downregulation in ECM-related genes and those found in the perineuronal nets of the synaptic cleft (TNC, THBS2) in SMA patient-derived motor neurons. We also observe an increase in several neurotransmitter receptors (SYNDIG1, GRM2, GRIK1), neurotransmission release and synaptic vesicle trafficking (RPH3A) that could be linked to SMA motor neuron hyperexcitability.

**Figure 4.**
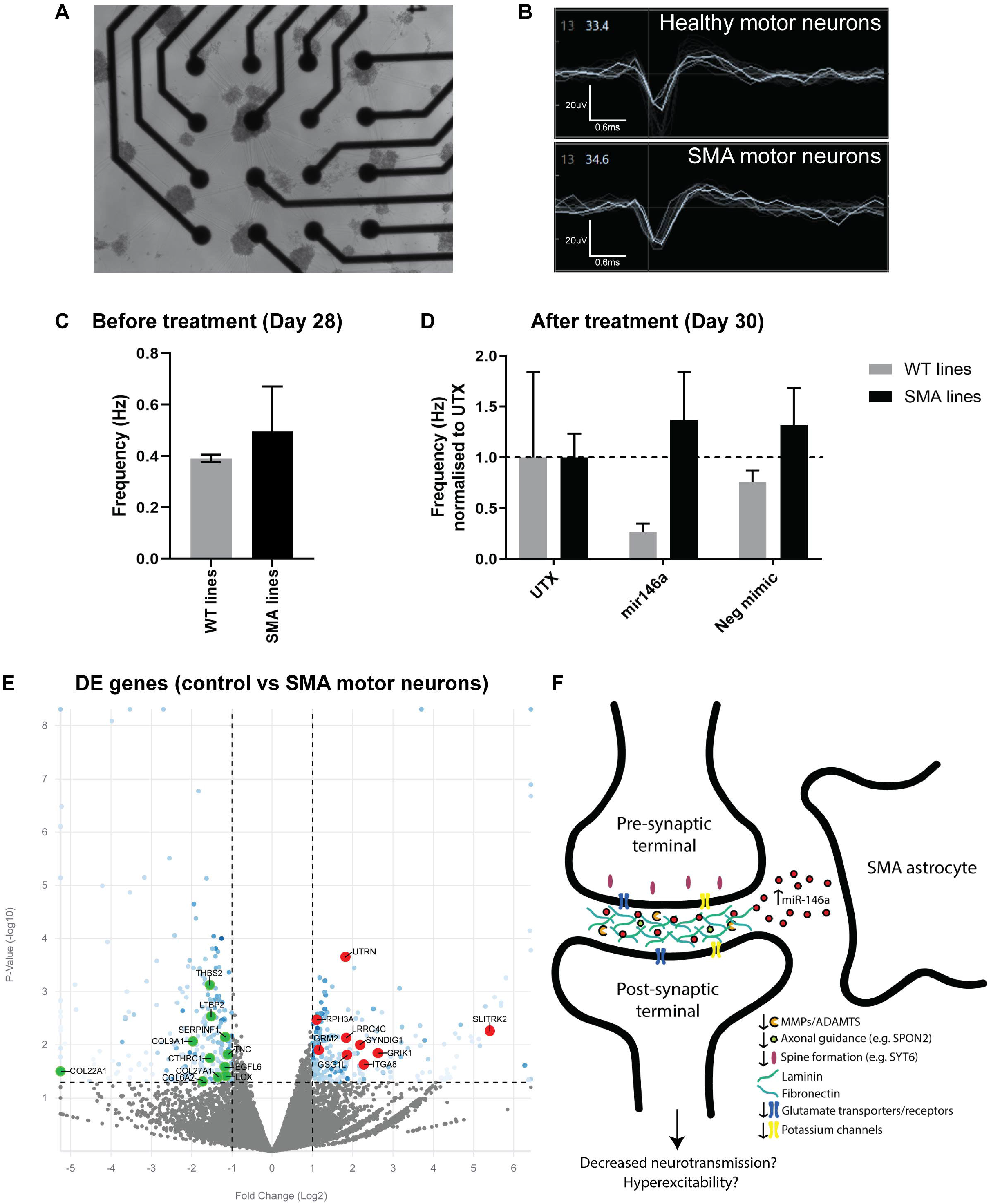
Multi-electrode array (MEA) analysis on miR-146a treated iPSC-derived motor neurons. To test the functional effects of miR-146a at the synapse, iPSC-derived motor neurons were plated on to the electrodes of MEA plates **(A)**. Actional potentials could be detected in both healthy and patient cells **(B)**. Weighed mean firing rate (WMFR) was measure before **(C)** and after **(D)** 48-hour treatment of either normal medium (UTX), 40nM miR-146a mimic or 40nM of neg miR mimic (neg mimic). Data for each line are normalized to WMFR for UTX sample. **(E)** DeSeq2 analysis reveals several ECM/ECM-synapse related genes that are downregulated in SMA motor neurons (green datapoints), whereas genes encoding synaptic-related genes are upregulated in SMA motor neuron (red datapoints). **(F)** Schematic of potential disease mechanism involving astrocyte secreted miR-146a at the perineuronal net of motor neuron synapses.

## Discussion

Despite the significant progress made in SMA therapy development, there remains a lack of reliably detectable biomarkers in patient biospecimens that can be used to track physiological change post-treatment over time. CSF is in direct contact with the primary diseased cells in SMA, and therefore represents a promising biospecimen to be used in biomarker discovery. Due to the intrathecal delivery of nusinersen, routine detection and analysis of biomarker trends in surplus CSF would be a feasible approach to perform in the clinic. For this study we focused on characterizing neurodegenerative biomarkers to provide a more disease-specific measurement to monitor CNS cellular health in SMA.

Clinically, all patients in this study receiving treatment did well, which is consistent to what has been seen in other patients treated with nusinersen and is contrary to the natural history of the disease (25, 26). Correlating these clinical observations to our biomarker screening, which overall showed continued upregulation of cellular stress genes but improvements in motor neuron-associated microRNA expression, further highlights the importance of understanding mechanisms driving cell recovery and survival and when improvements at the molecular level can be observed after nusinersen treatment. Our data also suggest that glial health is an important aspect to monitor in SMA patients in disease state and post-treatment, as demonstrated by the downregulated genes in the array and the continued elevated levels of astrocyte-secreted miR-146a in post nusinersen treated CSF samples. Post-mortem studies reveal that nusinersen is detectable glial cells throughout the CNS and spinal cord (27), but downstream molecular events after SMN targeting in human CNS cells have yet to be established. Our data suggest that longitudinal assessment of cellular function after prolonged nusinersen treatment (possibly greater than 13 doses; latest sample of this study) are needed beyond the motor function scores to fully elucidate changes in disease pathology.

While we found overlapping gene expression trends in the qRT-PCR array data for patients with the same disease severity, a number of genes were uniquely upregulated or downregulated in individual patient samples, and most of the genes analyzed did not show consistent expression trends across all patient samples (e.g. the same gene was upregulated in some samples, but downregulated in others). A similar observation was made in a proteomics study assessing neurodegenerative biomarkers in SMA CSF samples; no consistently differentially expressed proteins were found in response to nusinersen treatment across their dataset (15). The only consistent observation across all SMA patient samples post nusinersen treatment within our array data was the downregulation of ANGEL2, a recently identified CCR4 deadenylase with 2’,3’-cyclic phosphatase activity involved in RNA processing (23). ANGEL2 is thought to antagonize the mRNA splicing of XBP1, a transcription factor that regulates UPR during ER stress (28). Previous transcriptome data revealed SMA patient iPSC-derived motor neurons show upregulated expression of UPR genes, including spliced XBP1, compared to control iPSC-derived motor neurons (29). Downregulation of ANGEL2 within our dataset suggests cellular stress may still be apparent in patients even after nusinersen treatment. Future studies are needed to explore the potential mechanism involving ANGEL2 in SMA and its use as a biomarker in patient CSF samples.

Seems microRNAs are important RNA processing regulators, potentially linked to SMN function (30), can act as disease modifiers (31) and are relatively stable in biofluids, they represent good biomarker candidates. SMA patient serum has mainly been used to investigate microRNA levels at baseline (32) and after nusinersen treatment (33); seems microRNA specificity may differ depending on the tissue type (e.g. muscle vs CNS) and the type of biospecimen (e.g. serum vs CSF) analyzed, we focused on detecting microRNAs relevant to CNS cellular health in CSF samples. In most of the post-treated samples, we observed an overall increased expression trend of miR-9 and miR132, previously shown to be downregulated in SMA spinal cord tissue (32) and associated with neurite outgrowth delays (34-36). The same increased expression was also observed for miR-218 and miR23a normally downregulated in neuromuscular disorders, correlating with motor neuron synaptic defects and hyperexcitability (37), and motor neuron survival and neuroprotection (38). Our data are concordant with a previous study showing the improved increased expression of microRNAs (e.g. miR-132) in the spinal cords of mild and severe SMA mice after receiving PMO25 antisense oligonucleotide treatment (32). Together, this supports the notion that microRNA detection in CSF could serve as a useful way of monitoring physiological changes in response to treatment.

We have previously shown that SMA iPSC-derived astrocytes show abnormally elevated expression and secretion of miR-146a, which is sufficient to induce a decrease in the percent of ChAT+ motor neurons (19). The continued upregulation of miR-146a in CSF samples post nusinersen treatment prompted further investigation into mechanistic mediators via RNA seq, which could be involved in miR-146a induced motor neuron damage. Interestingly, the DeSeq2 analyses revealed a downregulation of genes related to extracellular matrix proteins, which complements findings from other human transcriptomic data comparing healthy and SMA iPSC-derived motor neurons (29). Many of the extracellular matrix associated genes within our dataset have been previously implicated at the synaptic perineuronal net, including HAPLN1 that was previously reported to be downregulated in SMA Type 1 and 3 patients (39), and those with altered expression in other neurodegenerative diseases, such as matrix metalloproteinases and NEDD9 (40, 41). Encouragingly, a number of these genes are validated miR-146a targets or have similar functional roles to known miR-146a targets, which implicates miR-146a-mediated regulation in synaptic function (42, 43). We also found a number of genes related to astrocyte mediated synaptic regulation, such as tenascin C and thrombospondin 2, which both play important roles at the perineuronal nets to support synaptic form and function (44). Additionally, evidence suggests that glial microRNAs can regulate ECM molecules, such as brevican (45), a major component of perineuronal nets. Therefore, we propose that miR-146a may be causing downregulation of synaptic related ECM genes and ultimately affecting perineuronal net regulation at the SMA motor neuron synapses.

Using a MEA approach, we also investigated the functional impact of miR-146a on synaptic activity of iPSC-derived motor neurons. Although we did observe a level of variability within the experiments, overall miR-146a treatment decreased firing rate of healthy motor neurons, whereas it increased activity in the SMA motor neurons. This was an intriguing finding in relation to the hyperexcitability phenotype previously identified in SMA motor neurons; evidence suggests this hyperexcitability phenotype could be a SMN-dependent motor neuron cell-autonomous (46) or non-cell autonomous phenotype (47). Our transcriptomic (healthy vs SMA samples; Fig. 4E) and MEA data suggest there may be a motor neuron intrinsic mechanism, seems we detect upregulated neurotransmitter receptor genes and increased baseline spontaneous action potential activity in SMA samples, in addition to observing opposite trends in miR-146a treated healthy and SMA-derived motor neuron WMFRs. However, together with the abnormally elevated miR-146a secretion by SMA astrocytes (19), the miR-146a transcriptome data also suggest a role of astrocyte-secreted factors causing dysfunction at motor neuron synapses in SMA. The exact pathways and downstream events that involve miR-146a and downregulated expression of these genes in SMA remains to be fully established, but our data begin to identify possibilities. For example, direct targets of miR-146a regulation, as previously demonstrated for a number of pre- and post-synaptic terminal genes (SYT1, NLG1) and those involved in synaptic plasticity (MAP1B) and glutamatergic neurotransmission (GRIA3, AMPARs, SLC17A8) may all contribute to motor neuron malfunction (42, 43, 48). The downregulation of neurotrophic factors (NTRK1, GFRA2) by miR-146a, could in turn affect the expression of motor neuron voltage-gated ion channels (49). A number of perineuronal net ECM associated genes downregulated in the miR146a treated motor neuron transcriptomes are known to play an active role in pre- and post-synaptic scaffolding (proteoglycans), synaptic stability and organization (cadherins, integrins), plasticity (tenascins), and synaptic remodeling (matrix metalloproteinases) to influence neurotransmitter receptor and ion channel activity (50). Therefore, there could be mechanistic links that involve ECM composition at perineuronal nets and regulation of their genes by astrocyte secreted miR-146a, ultimately impacting motor neuron expression of neurotransmitter and ion channel genes and leading to diminished functioning in SMA (Fig. 4F).

Together, our data demonstrate that SMA patient CSF exhibits altered transcriptional and glia-associated microRNA profiles indicative of sustained motor neuron stress and synaptic malfunction even with nusinersen treatment. Considering that treated SMA patients are now living long past the natural history of this disease, additional screening and profiling should be pursued to elucidate robust measures of therapeutic efficacy.

## Acknowledgements

The authors thank the patients and families for consenting to CSF sample collection and use. This study was supported by grants from CureSMA (ADE; EW).

## Author Contributions

Author contributions: E.W.: Acquisition and analysis of data, manuscript writing and editing. R.J.R.: Data acquisition. M.M.H.: Data acquisition and analysis. A.D.E.: Conception and design study, data analysis, manuscript writing and editing.

